# Nuclear size and physical properties of the nucleoplasm are determined by colloid osmotic pressure at the nuclear envelope

**DOI:** 10.64898/2026.07.21.739918

**Authors:** Joël Lemière, Zhidong Tan, Fred Chang

## Abstract

The size of the nucleus scales with cell size, suggesting a universal scaling rule. Yet the biophysical determinants of nuclear size and the significance and consequences of altered nuclear-to-cell (N/C) ratios, which are observed across diverse pathological states and cell-fate transitions, remain poorly understood. Recent theoretical models propose that nuclear size arises from a balance of colloid osmotic pressures generated by macromolecules in the nucleoplasm and cytoplasm. Here we demonstrate that altering this osmotic balance through massive overexpression of an exogenous protein targeted to either the nucleoplasm or cytoplasm produces predictable changes in the N/C ratio in *S. pombe*. These quantitative perturbations show that nuclear size is set primarily by the number of proteins in the nucleus and cytoplasm, providing strong support for a pure osmotic pressure mechanism. Furthermore, cells with altered N/C ratios display tunable changes in nucleoplasmic crowding, nuclear condensate formation, nucleolar scaling and heterochromatin organization, establishing a causal link between nuclear size and gene regulatory processes. These findings reveal how cells exploit osmotic forces to set organelle dimensions, with broad implications for understanding how nuclear size shapes gene expression and cell identity in health and disease.

## Results

### Addition of exogenous proteins into the nucleus or cytoplasm alters the N/C ratio

The observation that nucleus size scales with cell size — the so-called nuclear-to-cell volume ratio — has been recognized for over a century (Strasburger, 1893) and has motivated decades of study seeking to identify the underlying mechanism (Conklin, 1912; Hertwig, 1903; Harding and Feldherr, 1959; Neumann and Nurse, 2007; Moore et al., 2019; Cantwell and Dey, 2022). Yet only recently have physical models begun to formalize this relationship in quantitative, mechanistic terms. Recent studies propose that nuclear size is an emergent property arising from fundamental physical principles (Mitchison, 2019; Deviri and Safran, 2022; Lemière et al., 2022; Rollin et al., 2023). In this view, at steady state, nuclear volume is determined by a balance between colloid osmotic pressures generated by macromolecules in the nucleoplasm and cytoplasm, and membrane tension on the nuclear envelope. In contrast to cellular hydrostatic (or turgor) pressure that is generated by ions and small metabolites, colloid osmotic pressures are generated by collections of macromolecules such as proteins and nucleic acids. With the nuclear envelope acting a semi-permeable barrier that allows ions and small molecules to freely pass but not larger macromolecules (Mohr et al., 2009), the cytoplasm and nucleoplasm each harbor collections of macromolecules that generate their own colloid osmotic pressure.

A rigorous test of this osmotic model is to directly manipulate the number and subcellular localization of osmotically active macromolecules and assess whether nuclear size follows the model’s predictions. The model posits that addition of osmotically active proteins to the nucleus would increase relative colloid osmotic pressure in the nucleus, driving NE expansion, thereby increasing N/C ratio. Conversely, addition of proteins exclusively to the cytoplasm would increase the relative colloid osmotic pressure in the cytoplasm, leading to a decrease in the N/C ratio. Addition of proteins to both compartments would not be expected to affect the N/C ratio (**Fig. 1a**). Theoretically, the targeted expression of any osmotically active protein that is large enough so it cannot freely diffuse through the nuclear pores would have this property. However, to induce detectable changes in the N/C ratio, these proteins would need to be expressed at very high levels, representing a significant portion of the proteome.

**Figure 1.**
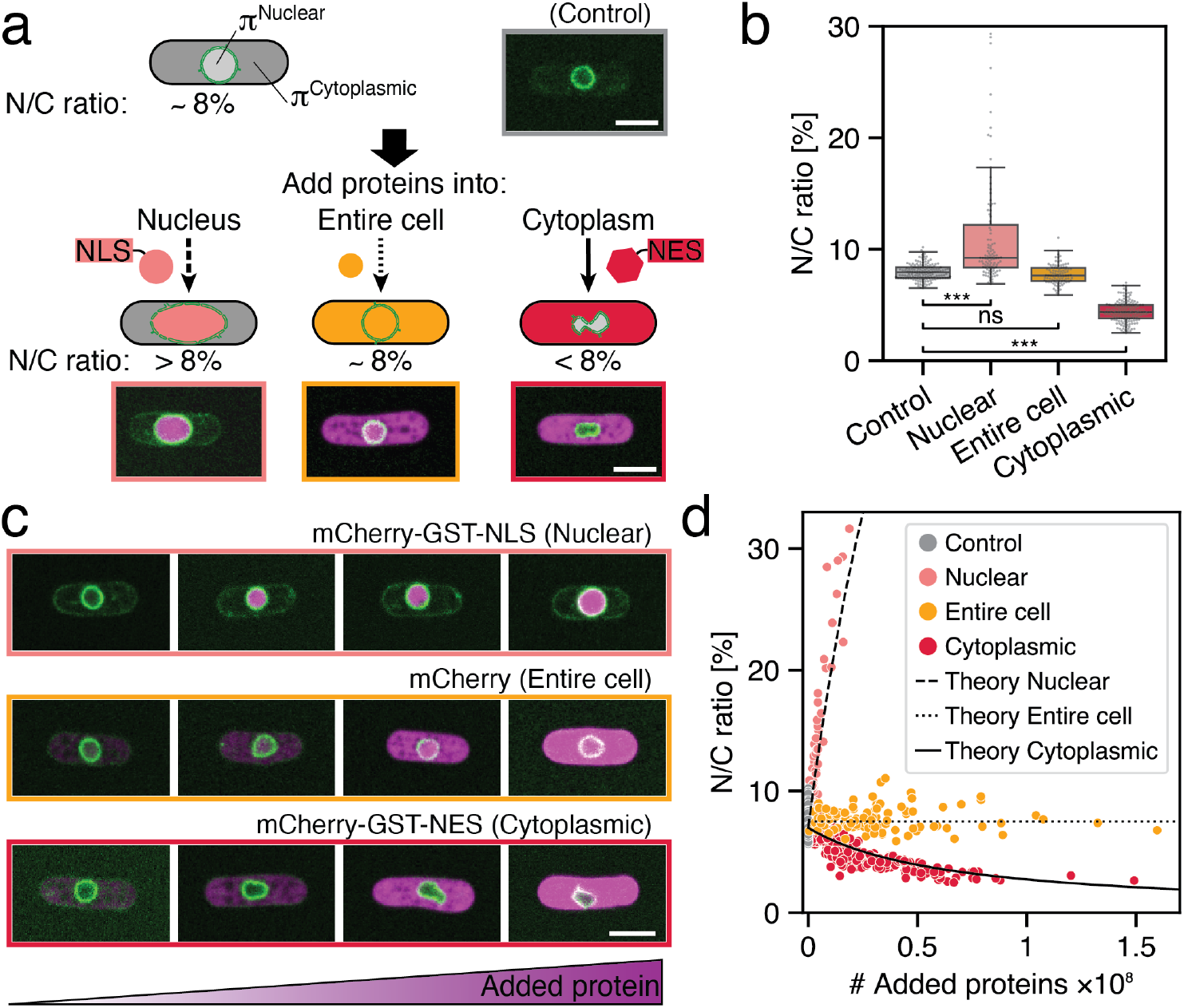
Addition of exogenous protein to the nucleus or the cytoplasm tunes the N/C ratio. **a,** Schematic illustrating the addition of nuclear and/or cytoplasmic proteins in fission yeast and the predicted effects on the N/C ratio. Inserts are representative middle plane confocal images of fission yeast expressing a nuclear envelope marker (Ish1-GFP, green) and mCherry (magenta) localized to the nucleus (mCherry-GST-NLS, left row), throughout the entire cell (mCherry, middle row), or to the cytoplasm (mCherry-GST-NES, right row). **b,** Quantification of the N/C ratio for populations of cells expressing the indicated protein constructs. N≥ 112 cells per conditions, two-sided Welch’s t-tests. **c,** Single-plane confocal images as in **a** (Nuclear, top; Entire cell, middle; Cytoplasmic, bottom), with exogenous protein expression increasing from left to right, revealing progressive effects on nuclear volume. Within each row, cells with similar volumes were selected to allow direct visual comparison of the nuclear size. **d,** N/C ratio as a function of the amount and subcellular localization of added exogenous protein (N=970 cells), overlaid with theoretical predictions of a pure osmotic model based upon protein number (see Methods). Statistical significance is indicated using standard asterisk notation: P ≥ 0.05 (ns), P < 0.05 (*), P < 0.01 (**), P < 0.001 (***). Scale bar, 5µm.

To achieve high-level expression of exogenous proteins with controlled subcellular localization, we engineered a series of constructs (**Supplementary Fig. 1a**). As mCherry can be expressed at very high levels in budding yeast (Kafri et al., 2016), we initially focused on mCherry-based fusions. Proteins were expressed in *S. pombe* from a strong promoter (P*tdh1*) using multi-copy plasmids based on pTOWsp-M backbone, which can amplify to up to 200 copies per cell (Moriya et al., 2011). Instead of mCherry alone, which is small enough (∼28 kDa) to diffuse passively through the nuclear pore, we used a larger fusion protein of mCherry and GST (glutathione-S-transferase; ∼54 kDa combined) to prevent passive nuclear diffusion and repeated cycles of nuclear shuttling. This avoids the nuclear transport defects that can arise from targeted mCherry proteins overwhelming the active nuclear transport machinery (Fujimoto et al., 2026). To target proteins to the nucleus, mCherry-GST was tagged with a nuclear localization signal (NLS). To restrict proteins to the cytoplasm, mCherry-GST was tagged with a nuclear export signal (NES). To extract N/C ratios alongside protein intensities from imaging data, we developed an automated image analysis pipeline. Briefly, nuclear and cellular volumes were segmented from spinning disc confocal fluorescence images, and protein intensities were quantified within each compartment (**Supplementary Fig. 2,3**).

Accumulation of mCherry-GST-NLS in the nucleus produced cells with strikingly large, round or ovoid nuclei with increases of N/C ratios up to 4-fold higher than normal (**Fig. 1a,b, Supplementary Fig. 2**). Accumulation of mCherry-GST-NES in the cytoplasm produced cells with abnormally small nuclei, with N/C ratios up to 2-fold lower than normal, consistent with increased cytoplasmic osmotic pressure pushing into the nucleus. These nuclei also displayed abnormal, indented morphologies, often in a characteristic bi-lobed shape. As a control, mCherry expressed from the same vectors localized to both the nucleus and cytoplasm and caused no change in N/C ratio or nuclear morphology (**Fig. 1a,b**). Collectively, these data demonstrate that targeting the accumulation of proteins to the nucleoplasm or the cytoplasm alters the N/C ratio.

These effects of mCherry-GST-NLS and mCherry-GST-NES expression on the N/C ratio were heavily dose dependent over a wide range of expression levels (**Fig. 1c,d**). We quantified the absolute number of mCherry molecules in individual cells by measuring fluorescence intensities. As calibration, we compared the fluorescence intensities of the mCherry fusions proteins to those of cells expressing histone Hta1-mCherry from its endogenous chromosomal locus, whose absolute number is estimated from proteomic studies and nucleosome counts ((Carme et al., 2026; Marguerat et al., 2012), **Methods**). This calibration was further validated by comparison to highly abundant proteins Eno1 and Tdh1 (**Supplementary Fig. 3 and Methods**) (Marguerat et al., 2012). These analyses show that the various mCherry-GST or mCherry proteins were expressed on the order of 10^6^ to 10^7^ proteins per cell, expression levels equivalent or surpassing the most abundant endogenous proteins in the cell (**Supplementary Fig. 3e**).

To test whether the effects of mCherry alone or fused to a GST-NLS or GST-NES on the N/C ratio reflect toxic properties specific to these proteins, we analyzed a series of control constructs. Replacing mCherry with GFP, substituting GST with tandem mCherry repeats, and varying the promoter and vector all yielded qualitatively similar changes in the N/C ratio (**Supplementary Fig. 4**). Furthermore, control cells carrying only the multicopy vector showed no change in N/C ratio compared to the N/C ratio of wildtype strain, indicating that the DNA plasmids per se did not affect nuclear size (**Fig. 1b,d**).

Expression of these fluorescent protein-derived constructs had surprisingly little effect on the growth or viability of the cells. The proteins did not affect cell size or morphology (length and width) (**Supplementary Fig. 5a-c**). No protein aggregates were detected by fluorescence microscopy even at the highest expression levels. Bulk growth curve assays showed no differences in exponential growth rates (∼4.10^−5^ h^−1^) and no evidence of widespread cell cycle delays or death (**Supplementary Fig. 5d**). However, time lapse imaging of individual cells revealed that the subpopulation with the highest levels of mCherry-GST-NLS expression exhibited growth inhibition, suggesting that the accumulation of proteins in the nucleus perturbs nuclear processes (**Supplementary Fig. 5e,f**). Together, these findings suggested that these exogenous proteins alter nuclear size through effects of osmotic pressure (see below) rather than from toxicity of these specific proteins.

### Quantitative validation of an osmotic model for nuclear size

With these experimental data, we next quantitatively tested the osmotic nuclear size model and addressed outstanding issues of osmotic phenomena inside cells. One critical question is: what are the relevant osmotic solutes contributing to nuclear size? Classic osmotic theory assigns equal contribution to all solute particles, regardless of size or charge, but in the crowded cellular milieu it is unclear which macromolecular solutes are the predominant generators of colloid osmotic pressure (Mitchison, 2019; Rollin et al., 2023). Proposals include proteins, RNAs, large macromolecular complexes, collective effects of chromatin with their counterions, as well as large metabolites. Another open issue is the role of macromolecular crowding: do solute concentrations in the cytoplasm and nucleoplasm place the system in a dilute regime or a non-dilute regime in which crowding interactions yield disproportionately high pressures (Ellis and Minton, 2003; Mitchison, 2019; Vink, 1971)?

To analyze our N/C ratio data, we applied a simple model in which nuclear size is set by the steady-state balance between colloid osmotic pressure in the nucleus and cytoplasm and membrane tension of the nuclear envelope (Deviri and Safran, 2022; Lemière et al., 2022) (**see model description in Methods**). Boyle Van’t Hoff experiments demonstrated that the *S. pombe* nucleus behaves as an ideal osmometer, indicating that nuclear envelope membrane tension is negligible (*σ_Nucleus_* ≃ 0 *mN*/*m*) (Lemière et al., 2022). We also assumed that osmotic solutes operate in a dilute regime, in which osmotic pressure (Π) scales linearly with solute concentration (c) and each macromolecule contributes equally to the total pressure (following van’t Hoff’s law). Equivalently, in a virial expansion of osmotic pressure, Π = RT(c + A_2_c^2^ + A_3_c^3^ + …), we set all virial coefficients beyond the first order to zero (Scatchard, 1946; Vink, 1974). We assumed that the solute behaviors are colligative – that is that they depend only on the number of total particles rather than on particle identity, charge or other properties. This model predicts that the N/C ratio is determined by the ratio of the numbers of macromolecules in each compartment (N^nucleus^ and N^cell^, equation (1).

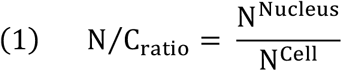

Addition of exogenous macromolecules to the nucleus (X) or cytoplasm (Y) lead to predictive changes in the N/C ratio as predicted by equations (2) and (3), respectively).

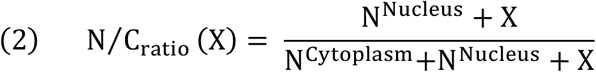

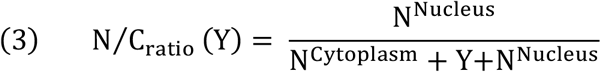

Plotting the theoretical predictions with our experimental data revealed a remarkably strong quantitative agreement without any free-fitting parameters (**Fig. 1d**). These experimental data therefore validate a pure osmotic model and support the assumption that osmolytes operate in a dilute colligative regime.

Using the nucleus as an osmometer, this quantitative framework allowed us to quantify colloid osmotic forces within the cell (**see Methods**). We showed that the nucleus is normally inflated by the osmotic pressure equivalent to 7×10^6^ mCherry-GST-NLS molecules in the nucleus, or approximately 5 mM in concentration; for example, the accumulation of 7×10^6^ mCherry-GST proteins into the nucleus resulted in approximately doubling of nuclear volume. This 5 mM concentration, using Van’t Hoff equation, corresponds to a colloid osmotic pressure of 12 kPa in the nucleus as well as cytoplasm within the unperturbed cell, similar in magnitude to previous estimates from proteomic data (Deviri and Safran, 2022) and representing ∼1% of the turgor pressure (Lemière and Chang, 2023).

Both mCherry-GST-NLS and mCherry-GST-NES data sets independently converged to an estimate of N^Cell^= 5±0.5×10^7^ osmotically active macromolecules per cell (see Methods). This number agreed remarkably closely with the total estimated number of proteins (6×10^7^) in a fission yeast cell based upon proteomic data (Schmidt et al., 2011; Marguerat et al., 2012; Wu et al., 2008). As further validation, we started with the published proteome numbers to show that model predictions closely fit with the experimental N/C results to both the Nuclear and Cytoplasmic (NLS and NES) data sets (**Supplementary Fig. 6a,b**). Independently, the mCherry molecule counts, which do not rely on the N/C ratio assumptions, lead to a prediction that 7.8% of the proteome resides in the nucleus, which would generate the N/C ratio of ∼7.5-8% under the pure osmotic model (**see Methods**). Thus, the osmotic behaviors of the nuclear and cytoplasmic proteomes are sufficient to explain quantitatively the determination of the N/C ratio.

The concordance between osmotically active solute counts and estimated protein number from proteomics indicates that most proteins in the collective proteome are osmotically active and account for most of intracellular colloid osmotic pressure. Together, these data support a pure osmotic colligative model in which the numbers of osmotically active proteins in each compartment (and not other solutes such as nucleic acids) produce the colloid osmotic pressures that set the N/C ratio.

### N/C ratio tunes the rheology of the nucleoplasm

One function of nuclear size control may be to regulate the concentration of components within the nucleus that determine physical properties of the nucleoplasm. To investigate how these perturbations impacts the physical properties of the nucleoplasm, we analyzed the diffusive-like motions of genetically encoded multimeric nanoparticles (GEMs) expressed from genomically integrated constructs (Delarue et al., 2018; Szoradi et al., 2021). GEMs targeted to the nucleus or cytoplasm enabled independent measurements of the nucleoplasmic and cytoplasmic rheology (**Fig. 2a, supplementary Fig. 7**). Particle tracks were analyzed by mean squared displacement (MSD) plots to derive effective diffusion coefficients (Deff) and the anomalous diffusion exponent (α). Expression of mCherry-GST-NLS in the nucleus led to a dose-dependent increase (up to 47% higher) in nuclear GEMs diffusivity, consistent with a less crowded, diluted nucleoplasm in the enlarged nucleus (**Fig. 2b,c**). In contrast, mCherry-GST-NES expression in the cytoplasm caused a dose dependent decrease in nuclear diffusivity in the shrunken nucleus, indicative of an increasingly dense nucleoplasm (**Fig. 2b,c**). Changes in the anomalous diffusion exponent showed consistent trends from a sub-diffusive behavior (α<1, reflecting hindered GEM movement) toward Brownian diffusion (α=1, indicating reduced confinement) with increasing N/C ratios (**Fig. 2c**). Accumulation of the mCherry in both compartments, which did not alter nuclear size, had no effect on nuclear GEMs diffusion coefficients or α values (**Fig. 2c**). The rheological changes were specific to the nucleoplasm, as none of the constructs affected GEMs diffusion within the cytoplasm (**Supplementary Fig. 7**). These impressive rheological changes correlated with the N/C ratio: diffusion increased in enlarged nuclei and decreased in small nuclei, with a four-fold range in the N/C ratio corresponding to a four-fold range in diffusion coefficients.

**Figure 2.**
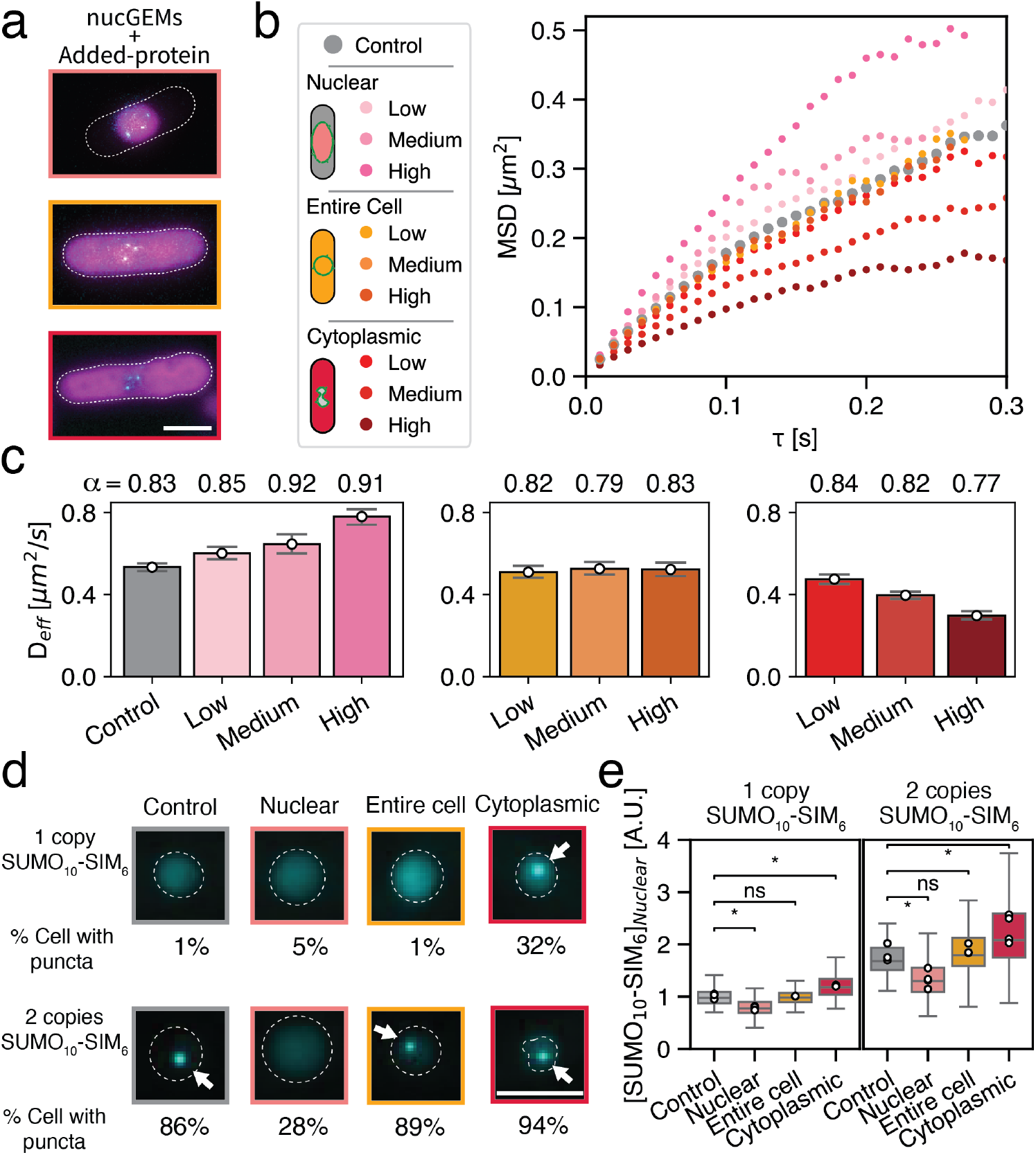
The changes in the N/C ratio alter the physical properties of the nucleoplasm. **a,** Representative fluorescence images of fission yeast expressing nuclear GEMs (cyan dots in nucleus) and the mCherry-based proteins (magenta). **b,** MSD plots of nuclear GEMs for each condition. Except for the control, the populations were split into low, medium, and high levels of mCherry expression. **c,** Effective diffusion coefficient (D_eff_) and the anomalous exponent (α) at various expression levels of the added protein. **d,** Cells expressing the synthetic condensate probe GFP-SUMO_10_-SIM_6_ (cyan). Representative summed Z-stack confocal images and the percentage of cells forming nuclear condensates for each condition (one copy, N = 608 cells and two copies, N = 534 cells). Dashed lines show nuclear outline, and arrow highlights nuclear condensate foci. **e**, Concentrations of GFP-SUMO_10_-SIM_6_ in the nucleus for each condition. Scale bar, 5 µm.

These rheological changes likely reflect global shifts in nucleoplasmic density and/or composition, driven in part by changes in nuclear volume. In cells with abnormally small nuclei due to cytoplasmic mCherry-GST-NES expression, the nucleoplasm loses water, resembling a hyperosmotic shock (Lemière and Chang, 2023; Molines et al., 2022), concentrating nuclear components without altering the cytoplasm. By contrast, in nuclei inflated by mCherry-GST-NLS, the exogenous protein may constitute up to 50% of the osmotically active nuclear proteome, while the endogenous proteins and DNA may be diluted by half because of the increase in the volume, assuming the absence of compensatory homeostatic mechanisms. Because osmotic pressure balance is maintained by changes in nuclear volume, the total concentration of osmotically active macromolecules in the nucleus may be unchanged in the enlarged nuclei. We propose that 40 nm GEMs diffuse faster because mCherry-GST protein, being much smaller than a GEM, is less potent as a crowding agent at that size scale than the typical nuclear macromolecules.

We next tested whether these perturbations to the rheology of the nucleoplasm affect the formation of liquid-liquid phase separation (LLPS) condensates within the nucleus. To assay for phase separation, we used a synthetic condensate probe that is based on tandem repeats of ten small ubiquitin-like modifier domains (SUMO_10_) and six repeats of SUMO-interacting motifs (SIM_6_) tagged with GFP and an NLS (Banani et al., 2016; Delarue et al., 2018; Sang et al., 2022). The SUMO_10_-SIM_6_ protein was expressed at two levels. (**Supplementary Fig. 8, 9**). At a low expression level (one copy of the SUMO_10_-SIM_6_ construct), GFP-labeled nuclear foci formed in mCherry-GST-NES cells (small nuclei), but not in controls or mCherry-GST-NLS cells (enlarged nuclei). At an approximately two-fold higher expression level (two copies of the SUMO_10_-SIM_6_ construct, **Supplementary Fig.8j**), foci appeared in ∼90% of the control and mCH-GST-NES expressing cells, whereas this frequency dropped to only 20% in the mCH-GST-NLS strains (**Fig. 2d, Supplementary Fig.8h and 9h**). Thus, these findings established an inverse relationship between the N/C ratio and the propensity for condensation within the nucleus.

The formation of condensate droplets is dependent on the concentrations of the phase-separating component(s) as well as mesoscale crowding factors (Banani et al., 2016; Alberti and Dormann, 2019, 2019; Shu et al., 2024). Fluorescence intensity measurements showed that for each series, SUMO_10_-SIM_6_ was expressed at the same levels per cell (**Supplementary Fig. 8f, 9f**), but its concentration in the different sized-nuclei varied inversely with nuclear size in a proportionate manner (**Fig. 2e, Supplementary Fig. 8g, 9g**). As indicated by the GEMs results, the N/C ratio may also affect the levels of macromolecular crowding in the different sized nuclei. We note that the addition of the exogenous mCherry-based proteins into the nucleus may only make a minor contribution to macromolecular crowing. For example, the mCherry control (Entire Cell) demonstrates that introducing an even large number of these exogenous proteins into the nucleoplasm itself does not affect diffusivity or phase separation if the N/C ratio is not altered. Thus, our findings highlight at least two ways in which changes in the N/C ratio mediates the tuning of the rheology of the nucleoplasm.

### N/C ratio tunes heterochromatin condensates

Given the observed effects on the synthetic condensate, we investigated whether expression of these exogenous proteins affect endogenous nuclear condensates. It has long been appreciated that in animal cells, increased nuclear volumes (or N/C ratios) are associated with decreases in heterochromatin and increased gene expression (Nicetto et al., 2019; Padeken et al., 2022; Shimada et al., 2009; Zhang et al., 2024). We thus tested a hypothesis that nuclear size may dictate the formation or maintenance of heterochromatin condensates. The heterochromatin factor Swi6 (an HP1 orthologue) (Grewal, 2023), which mediates heterochromatin nucleation and spreading in a dose-dependent manner, is one of the best characterized nuclear condensate proteins in fission yeast (Sadaie et al., 2008; Sanulli et al., 2019). It forms multiple nuclear foci that represent regions of heterochromatin at pericentromeric repeats, telomeres, and the mating-type loci near the inner nuclear membrane (Ekwall et al., 1995). Swi6 undergoes phase separation *in vitro*, in a manner dependent on its concentration and the concentration of crowding factors (Sanulli et al., 2019; Strom et al., 2017). *In vivo*, single molecule studies show that it dynamically exchanges between at least four pools: a mobile nucleoplasmic pool, a stable pool bound to methylated chromatin, a pool bound to DNA, and a condensate pool within chromatin representing approximately 20% of the total (Williams et al., 2024).

In the strains with a range of N/C ratios, Swi6-GFP foci were detected in all cases. As Swi6-GFP at the various nuclear loci normally display variable intensities, we first focused on comparing the intensities of just the centromeric foci (Kennedy et al., 2025) (the brightest spot in each cell, **see Methods**). The centromeric intensity changes were modest (less than 20%) but showed an inverse correlation with the N/C ratio: foci were dimmer in cells with enlarged nuclei and brighter in cells with small nuclei (**Fig. 3b**). A similar trend was observed in assessing all the types of Swi6 foci together (**Supplementary Fig. 10f**). In addition, the spacing of Swi6-GFP foci on the nuclear envelope correlated positively with the N/C ratio, with foci appearing closer together in smaller nuclei, demonstrating changes in chromosome organization with nuclear size (**Supplementary Fig. 10g**).

**Figure 3.**
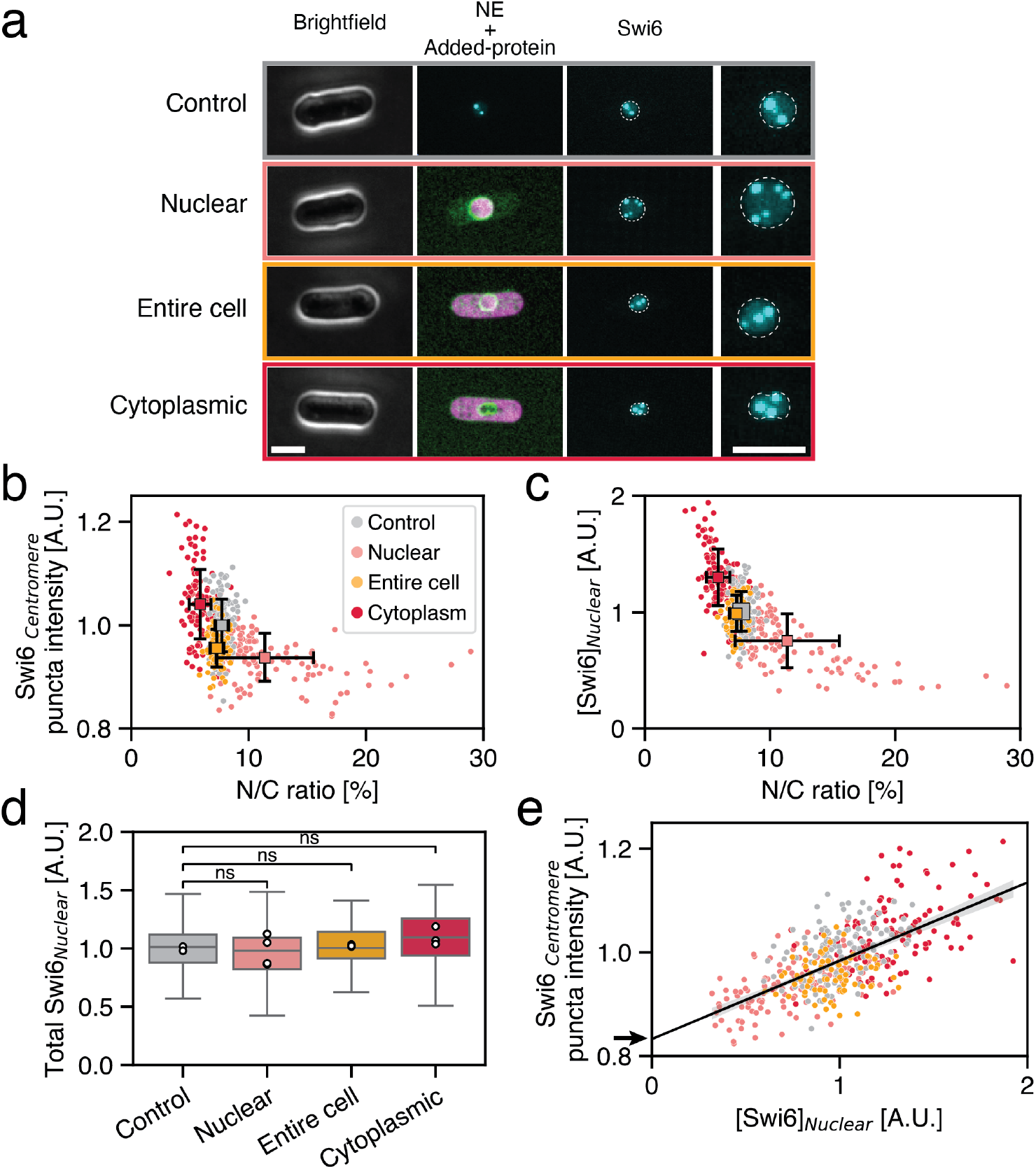
Alterations in the N/C ratio lead to changes in heterochromatin. **a**, Representative phase contrast and single-plane confocal images of fission yeast expressing various mCherry-based proteins (Fig.1a), with a heterochromatin marker Swi6-GFP (cyan) shown as a maximum projection. The rightmost column shows a magnified view of the nucleus, highlighted with a dashed line. Scale bar, 5 µm. **b**, Quantification of centromeric Swi6-GFP puncta intensity as a function of the N/C ratio (N = 512 puncta). **c**, Nuclear Swi6 concentration plotted as a function of the N/C ratio for individual cells, overlaid with mean values and standard deviations per condition (N = 1741 cells). **d**, Total nuclear Swi6-GFP intensity by condition, One-way ANOVA on replicate means. **e)** Centromeric Swi6-GFP puncta intensity as a function of nuclear Swi6 concentration, showing a linear positive correlation highlighted by the linear curve fit. The stable (non-condensate) fraction is determined by the y-intercept (black arrow). In b–c, 40% of the entire-cell condition data points were plotted for visualization.

As with the synthetic condensate, these nuclear size-dependent changes in Swi6 foci intensity may arise from changes in the concentrations of Swi6 and other heterochromatin factors in the nucleus. Consistent with this, Swi6 nuclear concentrations scaled inversely with the N/C ratio; for instance, a doubling of the N/C ratio led to an approximately two-fold decrease in Swi6 concentration and a 20% drop in Swi6 foci intensity at the centromeres (**Fig. 3c, Supplementary Fig. 10h**). Swi6 expression per cell was largely maintained across the range of N/C ratios, with no evidence of homeostasis regulation that maintains its concentration, and so Swi6 concentrations were directly regulated via nuclear size **(Fig. 3d**). Our data also revealed a linear positive relationship between Swi6 concentration in the nucleus and the intensity of Swi6 at centromeric foci (**Fig. 3e**). Assuming the phase separation model in which condensate fraction of the Swi6 puncta follows Lever rule (Alberti et al., 2019; Weber and Brangwynne, 2015), where the proportion of Swi6 partitioning into the condensate increases linearly with concentration, the slope predicts that ∼20% of Swi6 in the foci resides in a liquid condensate phase, while the y-intercept of 0.82 (**Fig. 3e, black arrow**) indicates that ∼80% is associated with chromatin in a concentration-independent manner (i.e. not a condensate). These observations match the estimates of the condensate pool obtained independently from single molecule dynamics (Williams et al., 2024). A similar positive correlation between Swi6 loci intensity and nuclear concentration was observed in assessing all Swi6 foci together (**Supplementary Fig. 10i**). These data suggest that changes in nuclear size leads to changes in Swi6 concentration that modulates the phase separation of heterochromatin condensates. Taken together, our findings demonstrate concentration-dependent mechanisms by which nuclear size modulates heterochromatin condensate states.

### Nucleolus size scales with cell size not nuclear size

Another prominent phase-separated organelle is the nucleolus, which serves as a ribosome biogenesis factory. The volume of the nucleolus has been shown to scale with nuclear or cell volume (Chen et al., 2024; Jorgensen et al., 2007; Neumann and Nurse, 2007; Weber and Brangwynne, 2015). However, as nuclear and cellular sizes are linked, it is not known if the principal scaling relationship of the nucleolus is to cellular volume or nuclear volume. Using Gar1-GFP, a nucleolar protein which is associated with small nucleolar RNAs, as a nucleolar marker (Girard et al., 1993), we measured nucleolar volume in the various strains (**Supplementary Fig. 11**). In wildtype cells with a normal N/C ratio, nucleolar volume scaled proportionally with nuclear volume with a volume ratio Nu/N=38±6 % (**Fig. 4a,b**) and also scaled with cell volume with a ratio of Nu/C= 3±1 % (**Fig. 4c**).

**Figure 4.**
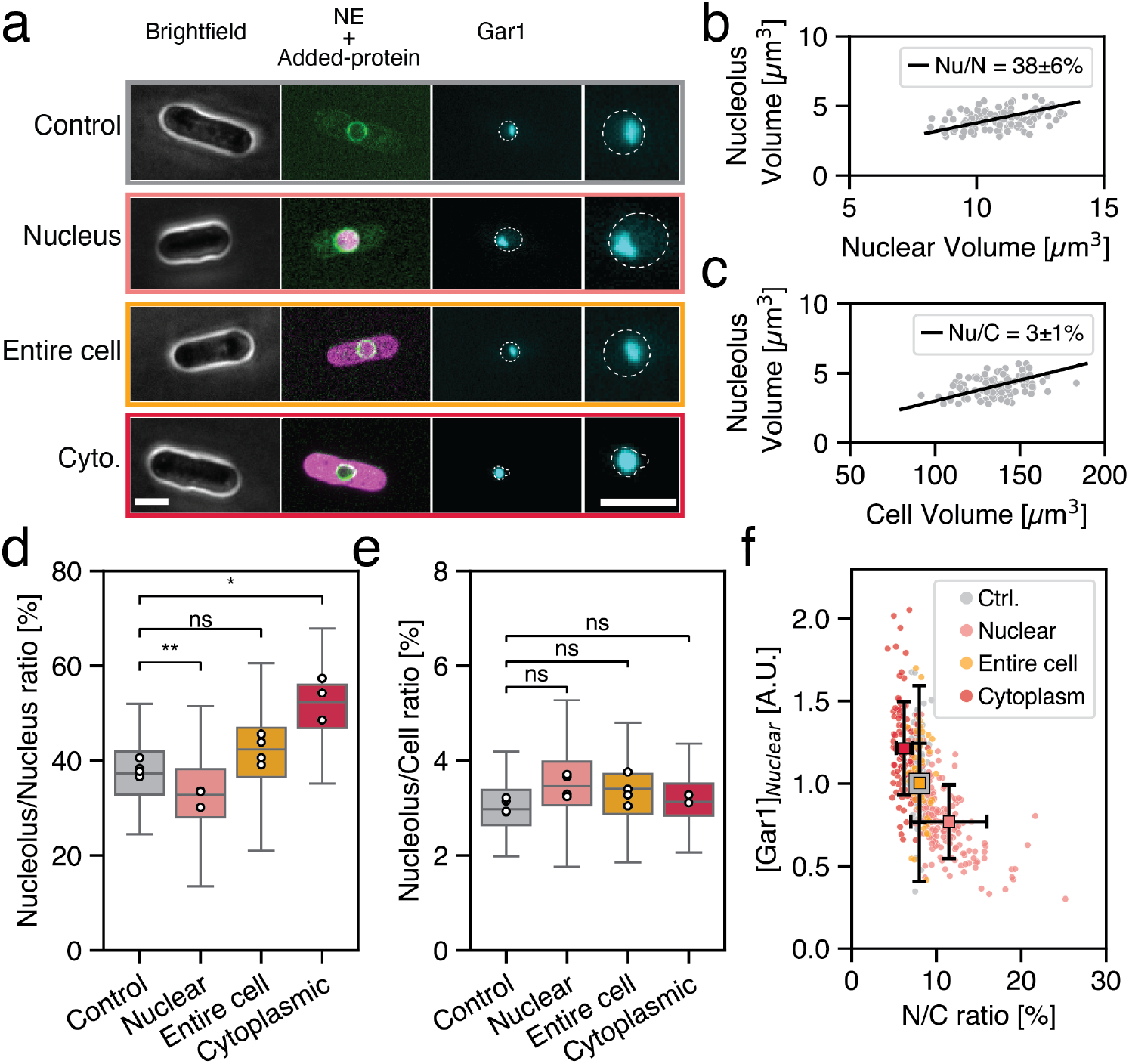
Alterations in the N/C ratio uncouple nucleolar volume from nuclear volume. **a,** Representative single plane confocal images of fission yeast cell expressing a nuclear envelope marker (green) and various exogenous mCherry-based proteins (magenta, Fig.1a), and sum projection images of a nucleolar marker Gar1-GFP (cyan) of the same cell, with a magnified view of the nucleus (right column) indicated by a dashed line. Scale bar, 5 µm. **b-c,** Scaling relationship of nucleolar volume with nuclear volume (b) and cellular volume (c) in control cells. The black lines represent a linear fit corresponding to a nucleolus to nucleus volume ratio Nu/N ∼38%, and nucleolus to cell volume ratio Nu/C ∼3% (N=104 cells). **d,** Nucleolus to nucleus volume ratio measured for each condition, showing a significant difference relative to the control cells only when mCherry expression is targeted to the nucleus or cytoplasm. (N≥97 cells per conditions). **e,** Nucleolus to cell volume ratio measured in the same cells as in (d) for each condition, showing no significative difference between conditions. One-way ANOVA statistical tests on replicates means. **f,** Nuclear Gar1-GFP concentration plotted as a function of the N/C ratio for individual cells, overlaid with mean values and standard deviations per condition (N=505 cells). Statistical analyses were performed on replicate means (white dots) using a one-way ANOVA, followed by pairwise Welch’s t-test comparing each strain to the control.

In the mCherry-GST expressing strains with altered N/C ratios, nucleoli were present but exhibited abnormal morphologies. In fission yeast, the nucleolus is normally a single, crescent-shaped structure located along a portion of the nuclear envelope (**Fig. 4a**). In cells with an enlarged nucleus, the nucleolus was not enlarged but showed decreased association with the nuclear envelope and was rounded rather than crescent-shaped (**Fig. 4a**). In cells with a shrunken nucleus, the nucleolus formed a rounded shape that takes up about half of the nuclear volume (**Fig. 4a**). In these cells, the nucleus was often a bilobed structure with one of the lobes filled with a rounded nucleolus, and the other with the rest of the nuclear components (**Fig. 4a**). Control cells expressing mCherry without changes in nuclear size exhibited nucleoli with normal morphology.

Volume measurements showed that in cells with a range of nuclear sizes, the nucleolar/nucleus ratios were significantly different but that nucleolar/cell ratios were maintained (**Fig. 4d,e**). The overall expression level of Gar1 was constant across conditions (**Supplementary Fig. 11f**), so that its concentration was inversely proportional to the N/C ratio (**Fig. 4f**). These data demonstrate that the nucleolus sizes scales with cell size (or cytoplasmic volume) rather than nuclear size. These findings are consistent with the proposal that nucleolar size is a barometer of the biosynthetic capacity of the cell and therefore may scale with the number of ribosomes in the cytoplasm (Tiku et al., 2017; Uppaluri et al., 2016).

## Discussion

These studies establish a systematic approach to experimentally manipulate nuclear size and the mechanical properties of the nucleoplasm in a defined manner. Tuning intracellular colloid pressures through massive expression of a single osmotically active protein in either the nucleoplasm or cytoplasm produces quantitatively predictable changes in the nuclear-to-cell volume ratio. These findings provide the most direct experimental validation to date of the colloid osmotic pressure model of nuclear size (Lemière et al., 2022; Mitchison, 2019; Rollin et al., 2023). Two independent perturbations (nuclear and cytoplasmic accumulation of exogenous protein) converge on the same estimate of total osmotically active macromolecules per cell (∼5 × 10^7^ or 5 mM), which corresponds to a colloid osmotic pressure of 12 kPa, roughly 1% of the ∼ 1 MPa turgor pressure in *S. pombe* (Lemière and Chang, 2023). The remarkable close agreement of this number of solutes with the number of proteins estimated from proteomic data implies that nuclear size is dictated primarily by the spatial distribution of the proteome, rather than nucleic acids, counterions or other macromolecular solutes. Our data further demonstrate that the proteins act in near-dilute, colligative regime at physiological concentration, which addresses a long-standing question in osmotic theory (Mitchison, 2019).

The ability to alter N/C ratios allows us to define the impact of nuclear size on cellular functions. Our findings demonstrate how the N/C ratio dictates the physical properties of the nucleoplasm in terms of diffusivity and crowding, simply by altering nuclear volume. Changes in the concentration of nuclear components lead to dose-dependent processes such as heterochromatin formation and others based upon liquid-liquid phase separation. Our studies provide insights into how nuclear size may contributes to changes in chromatin regulation and gene expression through the assembly or disassembly of heterochromatin factors. Cells may encounter analogous changes in the N/C ratio and the composition of the nucleoplasm and cytoplasm in developmental programs, aging and disease states (Jevtić and Levy, 2015; Neurohr et al., 2019; Syed et al., 2021; Zink et al., 2004; Zwerger et al., 2011). For example, expression of viral proteins during a viral infection (Herzog et al., 2026; Monier et al., 2000) or chromatin decondensing during NETosis, a form of neutrophil cell death (Thiam et al., 2020) may drive pathological changes in nuclear size. These studies therefore provide a novel framework to engineer nucleoplasmic properties that alter not only nuclear size but also cellular fate and disease states.

### Limitations of this study

Although we have employed controls to address these concerns, it is difficult to entirely rule out unintended effects of the artificial solutes such as mCherry-GST on cellular physiology, for example through off-target interactions with endogenous components, and indirect effects on transport, stress or biosynthesis. We argue however that the fit with quantitative predictions, consistent trends over a large range of increased and decreased N/C ratios, and the generally robust growth of these cells provide strong support for our biophysical interpretations.

The striking agreement of our mCherry solute counts and the estimated total proteins number support a pure osmotic model based upon macromolecular number, but it is possible that this concurrence in numbers is fortuitous. Protein count estimates carry appreciable uncertainty even for a given method (Ho et al., 2018). Moreover, not all endogenous proteins may be osmotically active because they are stably sequestered in large complexes such as chromatin (Mitra et al., 2025; Rao et al., 2007; Thiam et al., 2020). The relevant counts of mCherry-GST molecules themselves could also be an overestimate, as these proteins may form dimers due to the dimerization activity of GST (Fabrini et al., 2009). These studies also do not rule out alternative osmotic models based upon charges of macromolecules and their counterions, as described in pump-leak models that integrate the charge balances (Rollin et al., 2023). Future studies on effects of varying protein charge and assembly states will enable us to further refine various osmotic models.

Finally, these studies focus on nuclear size control in fission yeast, where osmotic forces predominate over nuclear envelope tension effects (Lemière et al., 2022; Oswald et al., 2006). Nuclear size in mammalian cells is also osmotically controlled but is partially constrained by membrane tension from the nuclear envelope (Finan et al., 2009; Oswald et al., 2006), which may arise from nuclear lamina, perinuclear actin and/or chromatin interactions (Marin et al., 2024). Future studies will be needed to integrate these additional factors to develop a full understanding of nuclear size control in more complex systems.

## Materials and Methods

### Yeast strains and media

The *Schizosaccharomyces pombe* strains used in this study are listed in **Supplemental Table S1**. In general, fission yeast cells carrying plasmids were grown at 30 °C in EMM3S – Edinburgh Minimum Media (#4110-32, MP Biomedicals) supplemented with 0.225 g/L of leucine, histidine, and adenine (#U0750, #H8000, #A9126, Sigma-Aldrich).

For stable expression of cytGEM and nucGEM proteins at the population level, we integrated the Pyrococcus furiosus encapsuling gene encoding 40 nm-GEMs fused to a fluorescent protein, mSapphire, into the *ura4+* locus using standard PCR-based methods (Bähler et al., 1998). For nuclear GEM probes (nucGEMs), we used the pRga3 promoter to drive expression. For cytoplasmic expression (cytGEMs), we used the intermediate strength pHis3 promoter (Carpy et al., 2014; Vještica et al., 2020).

### Microscopy

Cells were imaged on a Ti-Eclipse inverted microscope (Nikon Instruments) with a spinning-disk confocal system (Yokogawa CSU-10) that includes 488 nm and 541 nm laser illumination and emission filters 525 ± 25 nm and 600 ± 25 nm respectively, a Borealis illumination system (Andor) for even illumination, a 60X (NA: 1.4) objective, and an EM-CCD camera (Hamamatsu, C9100-13). These components were controlled with µManager v. 1.41 (Edelstein et al., 2010, 2014). Temperature was maintained at 30°C by a black panel cage incubation system (#748-3040, OkoLab).

For confocal live cell imaging, cells were mounted on a 1% agarose (Invitrogen, #16500500) pad sealed with VALAP and incubated for 15 minutes before imaging.

For imaging of GEMs, live cells were imaged with a TIRF Diskovery system (Andor) with a Ti-Eclipse inverted microscope stand (Nikon Instruments), 488 nm laser illumination, a 60X TIRF oil objective (NA:1.49, oil DIC N2) (#MRD01691, Nikon), and an EM-CCD camera (Ixon Ultra 888, Andor), controlled with µManager v. 1.41(Edelstein et al., 2010, 2014). Temperature was maintained at 30°C by a black panel cage incubation system (#748-3040, OkoLab). Cells were mounted in µ-Slide VI 0.4 channel slides (#80606, Ibidi – 6 channels slide, channel height 0.4 mm, length 17 mm, and width 3.8 mm, tissue culture treated and sterilized). The µ-Slide channel was coated by pre-incubation with 100 µg/mL of lectin (#L1395, Sigma) for 15 min at room temperature and then washed with medium. Cells in liquid culture were introduced into the chamber, incubated for 5 min and then washed three times with medium to remove non-adhered cells.

### 3D volume measurements

Nuclear and cell volumes were measured in individual cells using a pipeline as described (**Supplementary Fig. 2**). *S. pombe* strains expressing a nuclear envelope marker (Ish1-GFP, Ish1-mCherry or Ish1-CFP) and a protein of interest labeled with mCherry or GFP were imaged using phase-contrast and spinning disc confocal microscopy to acquire z-stacks (0.5 µm spaced). To define cell contours, the phase-contrast image of the middle plan was used for gradient segmentation at sub-pixel resolution in Morphometrics (Ursell et al., 2017).

Taking advantage of the cylindrical geometry of fission yeast cells, we apply principal component analysis to the contour to calculate two main symmetry axes, which approximate the cell’s length (long axis) and width (short axis). Cell length was determined through a step-by-step method: starting at the center of the contour and following the long axis, symmetric bisections transform the middle axis into a midline closely fitting the contour. Assuming rotational symmetry around this midline, cell volume is calculated by integrating frustums of cones along the midline. The center of the contour is assumed to indicate the nucleus’s position. This position is then applied to the fluorescent z-stack images to segment the nucleus in 3D using the ImageJ plugin LimeSeg (**Supplementary Fig. 2a**). Once the cell and nucleus were segmented, fluorescence quantification of the protein of interest was automated, enabling measurement of protein levels per cell in the nucleus and the cytoplasm.

This approach based on 2D segmentation of cellular volume was validated by comparing it to a pre-existing method that relies solely on 3D segmentation (**Supplementary Fig. 2b**)(Machado et al., 2019). Analyses of images of cells with labeled nuclear and plasma membranes showed a slight increase in the measured cellular volume using 2D segmentation compared to 3D segmentation across the entire cell population. The two methods showed good overall agreement (R^2^ = 0.87), however, 2D segmentation systematically overestimated cellular volume relative to 3D segmentation, with a mean offset of 2.70 µm^3^ and a mean absolute error of ∼6 % (**Supplementary Fig. 2c**). This small discrepancy in cellular volume was then carried over to the N/C ratio, explaining the slight (∼2%) increase in the 3D measurement compared to 2D ones (**Supplementary Fig. 2d**). Overall, these data demonstrate that this new pipeline, which combines 2D segmentation of cellular volume with 3D segmentation of nuclear volume, is in good agreement with previously published values (Lemière et al., 2022; Lemière and Chang, 2023). This automated segmentation pipeline allowed us to analyze 100’s of cells and perform time-lapse analyses to track individual cells over time.

To measure nucleolar volume, we imaged cells expressing the nucleolar protein Gar1-GFP expressed from its chromosomal locus (Carme et al., 2026; Girard et al., 1993). As the nucleolus was fully labeled with Gar1-GFP, the nucleolar volume was quantified based upon the number of bright voxels. The background fluorescence was first subtracted using the rolling-ball algorithm (radius = 25 pixels) applied to the entire z-stack to enhance signal contrast. An automatic threshold was then determined using the MaxEntropy method, and the images were converted into binary masks to isolate nucleolar voxels. The stack was subsequently projected using the “Sum Slices” Z-projection to obtain the total number of pixels corresponding to the nucleolus. The bright pixels were then calibrated using the voxel dimensions (pixel width, height, and Z-step) to convert the signal into a volume measurement in µm³.

### Fluorescent intensities quantification

mCherry fluorescence intensity was measured using defined ROIs. After cell and nucleus segmentation (**see 3D Measurement Method section**), the nuclear position (X_N_,Y_N_) was used as the center of the nuclear ROI (**Supplementary Fig. 2a**). For each half of the cell’s long axis, the midpoint between the nucleus and the cell tip was calculated to define two cytoplasmic ROIs (X_C1_,Y_C1_, and X_C2_,Y_C2,_ (**Supplementary Fig. 2a**)). Finally, a fourth ROI was automatically selected outside of the cell to account for background intensity (X_B_,Y_B_,, (**Supplementary Fig. 2a**)). From these coordinates, we automatically measured the mCherry fluorescence intensity each ROI in the middle plane of each z-stack. For the cytoplasmic signal, we averaged the intensities measured on each side of the cell. The background fluorescence signal was then subtracted from both the nuclear signal and cytoplasmic signals. To measure the total mCherry intensity for each compartment, the average background-corrected intensity per compartment was multiplied by its respective volume, either the nuclear or the cytoplasmic volume.

mCherry Intensity calibration and quantification: To calibrate the intensity of our measurements, we imaged a population of exponentially growing wild-type cells expressing a nuclear envelope marker (Ish1-GFP) and histone Hta1 tagged with mCherry (Hta1-mCherry, (Carme et al., 2026)) at its chromosomal locus. For each cell, the nuclear position was determined via 2D segmentation of the full-cell phase image (see 3D Measurement Method section). This ROI was then used to seed LimeSeg (Machado et al., 2019) and recover the nuclear volume. The average pixel intensity of Hta1 within this ROI was measured, assuming homogeneous distribution within the nucleus. The total tagged histone intensity for each cell was obtained by multiplying the average pixel intensity with the nuclear volume. To ensure consistency across experimental replicates, the mean Hta1-mCherry intensity measured across a population of cells was calculated for each experiment. The ratio between this mean and that of the first replicate was then used as a correction factor, which was applied to all fluorescence intensity measurements acquired on the same day. Finally, Hta1 intensities were converted into protein numbers using a calibration factor. The mean intensity across all calibrated replicates corresponded to the number of Hta1 molecules determined in a proteome quantification study of an exponentially growing population of *S. pombe* cells (N^Hta1^=1.4 ×10^5^, (Marguerat et al., 2012)). As a cross-check, a back-of-the-envelope calculation was performed to estimate the number of Hta1 molecules per cell, yielding a consistent result: assuming that each nucleosome wraps approximately 200 bp of DNA and contains two Hta1 molecules and given that the *S. pombe* genome is ∼12.6 Mbp, this yields an estimate of∼1.3 ×10^5^ Hta1 molecules per cell.

To validate our protein number quantification, we tagged two highly expressed proteins (Eno1 and Tdh1) with mCherry; both proteins had also been quantified in a previous quantitative proteomic study. Our values (N^Eno1^ = 3.1 ± 0.6 ×10^6^, N^Tdh1^ = 3.7 ± 0.9 ×10^6^) are within the same order of magnitude as published values (N^Eno1^ = 1.4 (0.6-3.4) × 10^6^, N^Tdh1^ = 1 (0.42-2.4) ×10^6^, (Marguerat et al., 2012)).

Nuclear protein quantification: Cells expressing GFP-tagged proteins from their native chromosomal loci, were imaged in Z-stacks with a 0.5 µm step size. For each field of view containing a single cell, we used the nuclear position obtained from our cell and volume measurement pipeline (see 3D Volume Measurement section) to define a square region of interest (ROI) centered on the nucleus in the sum-intensity projection of the GFP channel Z-stack. This ROI was used to measure the total nuclear fluorescence intensity of the GFP-tagged protein. A second ROI, located outside the nucleus, was automatically selected to measure background fluorescence, which was then subtracted from the nuclear signal. Total nuclear intensities measured from a population of control cells (containing the empty vector) were used to normalize intensities across days and replicates. The nuclear concentration of each protein was calculated by dividing the total nuclear intensity by the nuclear volume measured via our 3D volume measurement pipeline for each cell.

### Spectrophotometric growth curves

Cells were grown overnight in EMM3S at 30°C. Cultures were then diluted into fresh medium at an optical density at 600 (OD_600_) of 0.1, measured using a benchtop spectrophotometer (NanoPhotometer C40, IMPLEN). 300 µL of each culture was dispensed into a sterile multiwell plate (Nunclon Delta Surface, ThermoFisher scientific) and incubated at 30°C in a plate reader (Varioskan LUX, ThermoFisher Scientific). Absorbance of the cell suspension was recorded every 20 minutes with continuous shaking (540 rpm medium force setting) between readings. A well containing only EMM3S medium was included on each plate and used as a blank for calibration during the entire time of the experiments. For each condition (i) seven growth curves were averaged, and exponential (log-phase) portion of the resulting curves was fitted with a simple exponential function:

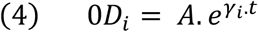

Where A is the initial OD_600_ at the start of the fitted log-phase and *γ_i_* is the population growth rate express in (h^−1^).

### Single cell growth curves

An initial Z-stack was acquired for each cell to enable quantification of their nuclear volume (ish1-CFP), and mCherry levels of expression (RFP). Ater this initial acquisition, phase contrast images were collected every 4 minutes for one hour. Using our cell segmentation tool, we measured growth as the increase in cellular volume in interphase cells. Growth curves were then normalized by the initial volume of each cell *V*(*t* = 0)*_i_* and fitted with a simple exponential to estimate single cell growth rate *γ_i_*:

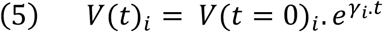

### Measurement of effective diffusivity and anomalous diffusion exponent

GEMs were tracked using Particle Tracker 2D/3D plugin from MosaicSuite in ImageJ (Sbalzarini and Koumoutsakos, 2005) with the following parameters: run(“Particle Tracker 2D/3D”, “radius = 3 cutoff = 0 per/abs = 0.03 link = 1 displacement = 6 dynamics = Brownian”).

The analyses of the GEMs tracks were described in (Delarue et al., 2018; Lemière et al., 2022; Molines et al., 2022), Mean square displacement (MSD) curves were computed using MATLAB (MATLAB_R2022, Math-Works). The effective diffusion D_eff_ was determined by fitting the first 10 time points of the MSD curve (MSD_truncated_) to the standard 2D Brownian motion equation: MSD_truncated_(τ) = 4D_eff_ τ. The anomalous exponent (*α*) was obtained by performing a linear fit of the logarithm of the MSD versus the logarithm of time over 100 ms interval.

For each condition, the average MSD and its standard deviation were computed as a function of time lag τ. To determine α, a linear regression was performed in log-log space on the first ten-time lag points, exploiting the power law relationship MSD ∼ τ^α^. The uncertainties on α represent the standard error of the fitted slope.

Multiple fields of view were acquired per condition. Individual cells were cropped from each field, and the mCherry fluorescence intensity channel was used to quantify expression levels. The full dataset was partitioned into three equal groups based on mCherry intensity, corresponding to low, medium, and high expression levels, for which D_eff_ and α were measured independently.

### Modeling the effect of adding osmotically active particles on the N/C ratio

The theory developed here is based on a balance of colloid osmotic forces at the nuclear envelope membrane. We assumed that the nuclear envelope membrane tension is negligeable (*σ_Nucleus_* ≃ 0 *mN*/*m*, (Lemière et al., 2022)), and that solute concentrations contribute linearly to the osmotic pressure (i.e., virial coefficients beyond first order are set to zero), such that the colloid osmotic pressure satisfies the Van’t Hoff equation P = cRT, where c is the concentration of solute particles, R is the ideal gas constant, and T is the temperature. The equilibrium of the colloid osmotic pressures between the cytoplasm and the nucleoplasm leads to the expression of the N/C ratio developed in our previous study (**Equation 6**),(Lemière et al., 2022)). If we now partition the total number of macromolecules within the cell (N^Cel^) between those in the nucleus (N^Nucleus^) and those in the cytoplasm (N^Cytoplasm^):

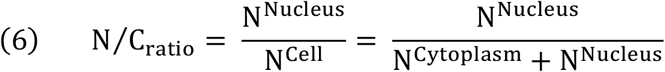

If we now assume that we can add osmotically active macromolecules specifically to the nucleoplasm or the cytoplasm (hereafter named X and Y, respectively), this equation can be written as:

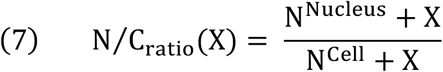

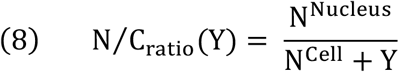

This assumes that every added protein is fully localized to either the nucleoplasm or the cytoplasm and contributes equivalently to the osmotic pressure, as captured by ideal gas theory.

To plot the model predictions (**Fig. 1d**), we assumed that the total amount of osmotically active macromolecules within the cell (N^Cell^) is equal to the total number of proteins in fission yeast. We took the value of N^Cell^ = 6.0×10^7^ from a quantitative proteomic analysis performed on a population of wildtype *S. pombe* in exponential growth phase as the total number of proteins per cell (Marguerat et al., 2012; Milo, 2013). Based on our assumption that what drives nuclear volume is the amount of protein within the nucleus, we estimated that 7.5% of this total amount resides in the nucleus, such that N^Nucleus^ = 4.5×10^6^. Finally, we assumed that the cell produced these extra proteins (X or Y) without changing the expression of the endogenous pool, so that the baseline nuclear and cytoplasmic protein counts are fixed. This allowed us to plot the predicted change in N/C ratio as a function of the quantity of exogenous proteins added to the nucleus (X) or the cytoplasm (Y) in **Fig. 1d**.

We also used our experimental data on altered N/C ratios as a function of the amount of exogenous protein added either to the nucleus (X) or the cytoplasm (Y) to estimate the number of osmotically active molecules within the cell and the nucleus. X and Y were calibrated for intensity against the amount of Hta1-mCherry protein as described in the previous paragraph. We then fitted these two curves, which allowed us to independently determine the total protein count and the nuclear protein count (i.e., when X = Y = 0). The fitted values yielded a total protein count of N^Cell^ = 4.4×10^7^ – 5.4×10^7^, which is in good agreement with the expected total protein count for fission yeast (6×10^7^) and with our assumption for the theoretical plots described in the first paragraph. These parameters and their uncertainties (1σ) were obtained by nonlinear least-squares fitting, with errors estimated from the covariance matrix of the fit (**Supplementary Fig. 6b**). These fits also provide an estimated nuclear osmolytes count of N^Nucleus^ = 3.2×10^6^ – 3.8 ×10^6^, likewise in good agreement with our model prediction (4.5×10^6^).

### Synthetic condensate formation

To probe phase-separation behavior in the nucleoplasm, we imaged a nuclear localized synthetic condensate probe based upon the fusion protein GFP-SUMO_10_-SIM_6_ (Delarue et al., 2018). We designed two variants of pAct1-NLS-GFP -10xSumo-6xSIM plasmids: we replaced the nuclear export signal (NES) present in pAV106-pTDH3-mCherry-10xSumo-6xSIM plasmid (Delarue et al., 2018) with an SV40 nuclear localization signal (NLS), and inserted the resulting constructs into the pAde6(PmeI)-p(act1)-sfGFP-term(ScCYC1) and pAV0663 pHis5StuI-p(act1)-sfGFP-term(ScCYC1) backbones, which provided the *S. pombe act1* promoter, sfGFP and the Ade6 or His5 selection cassettes, respectively (Vještica et al., 2020). We integrated a copy of the pAct1-NLS-sfGFP-10xSumo-6xSIM construct into the *ade6+* locus, while a second copy was subsequently inserted at the *his5*+ locus, yielding strains with one and two genomic copies of the construct, respectively. The two-copy strain showing 1.8 times higher expression level (**Supplementary Fig. 8h**). The expression of the SUMO_10_-SIM_6_ proteins did not alter the N/C ratio phenotypes (**Supplementary Fig. 8i**). Quantification of the nuclear SUMO_10_-SIM_6_ was done following the protocol described above (**Section Nuclear protein quantification**).

### Heterochromatin localization

For each field of view containing a single cell, we used the nuclear position obtained from our cell and volume measurement pipeline (see 3D Volume measurement section) to define a square region of interest (ROI) centered on the nucleus in the sum-intensity projection of the Swi6-GFP Z-stack. This ROI was used to measure the total nuclear intensity of Swi6-GFP fluorescence. A second ROI, located outside the nucleus was automatically selected to measure background fluorescence, which was subtracted from the nuclear signal.

Swi6-GFP foci were detected using TrackMate to determine their 3D positions (X,Y,Z) and fluorescence intensities. We used the LoG detector with an estimate object diameter of 0.8 µm, with subpixel localization enabled; and pre-processed with a median filter. The centromeric Swi6-GFP was defined as the brightest Swi6 punctum in each nucleus, which has been shown to colocalize with the Spindle Pole Body (SPB) protein Sad1 (Kennedy et al., 2025). A custom script on Jupiter Notebook in Python was used to automatically quantify, for each nucleus, the number of detected puncta and their distances to the SPB for each nucleus. Swi6–GFP nuclear concentration was computed using the method described in the section nuclear protein quantification.

## Acknowledgments

We are grateful to members of the Fred Chang, Sophie Dumont, and Orion Weiner labs for helpful discussions, and to Matthieu Piel for early conversations that helped initiate this project. We thank Gant Luxton for comments on the manuscript and Sophie Martin for providing plasmids. Plasmids FYP2790, FYP2791, and FYP2792 were provided by the National BioResource Project (NBRP) – Yeast, Japan. This project was supported by grants NSF 2213583 (to FC) and NIH R35 GM141796 (to FC).

## Author Contributions

Conceptualization: J.L. and F.C.; Methodology: J.L. and F.C.; Software: J.L.; Validation: J.L., Z.T.; Formal analysis: J.L.; Investigations: J.L. Z.T.; Writing: J.L. and F.C.; Review editing: J.L., F.C. Visualization: J.L.; Supervision: J.L., F.C.; Project administration and funding acquisition: F.C.

## Supplementary Material

**Table S1.**
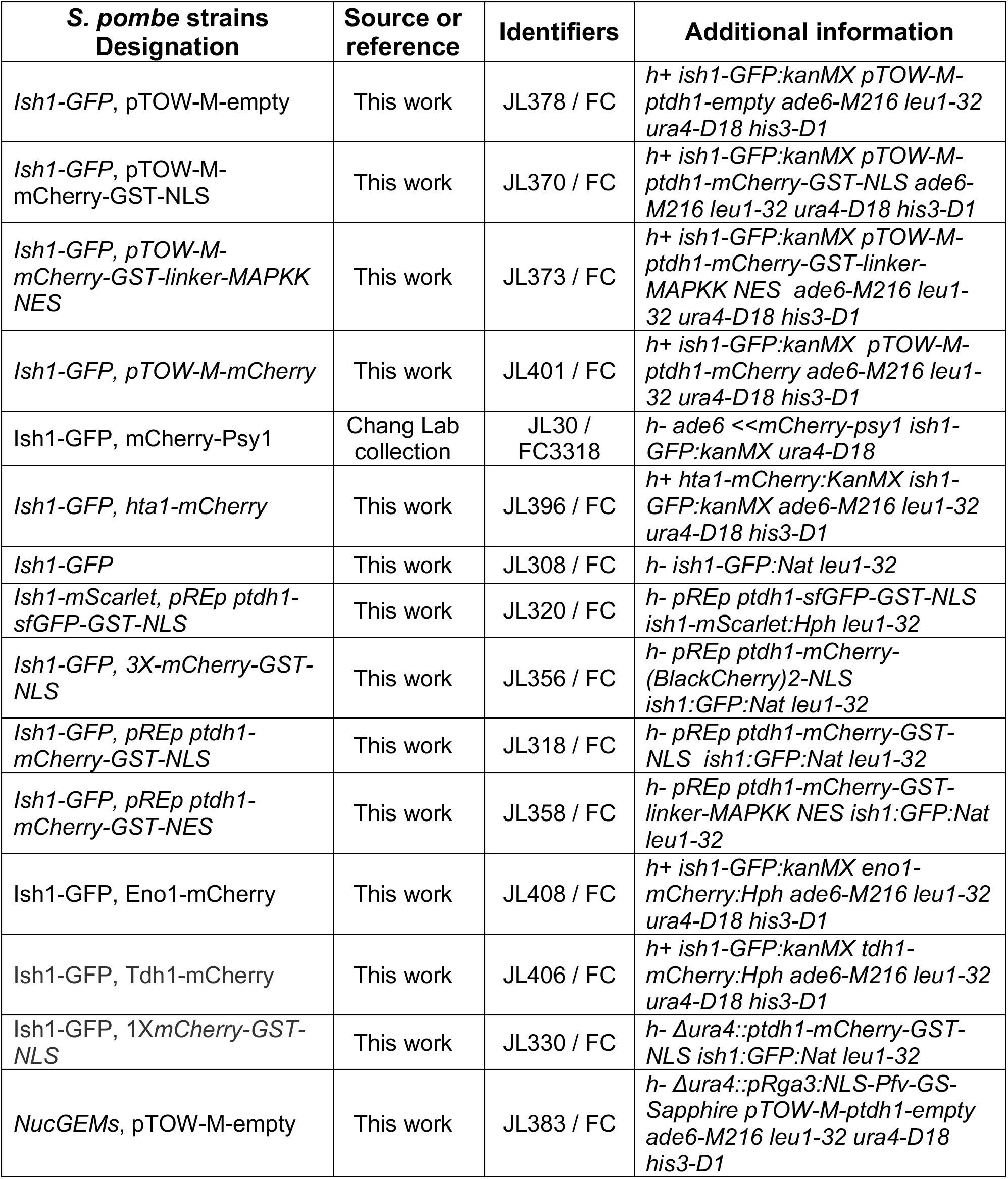
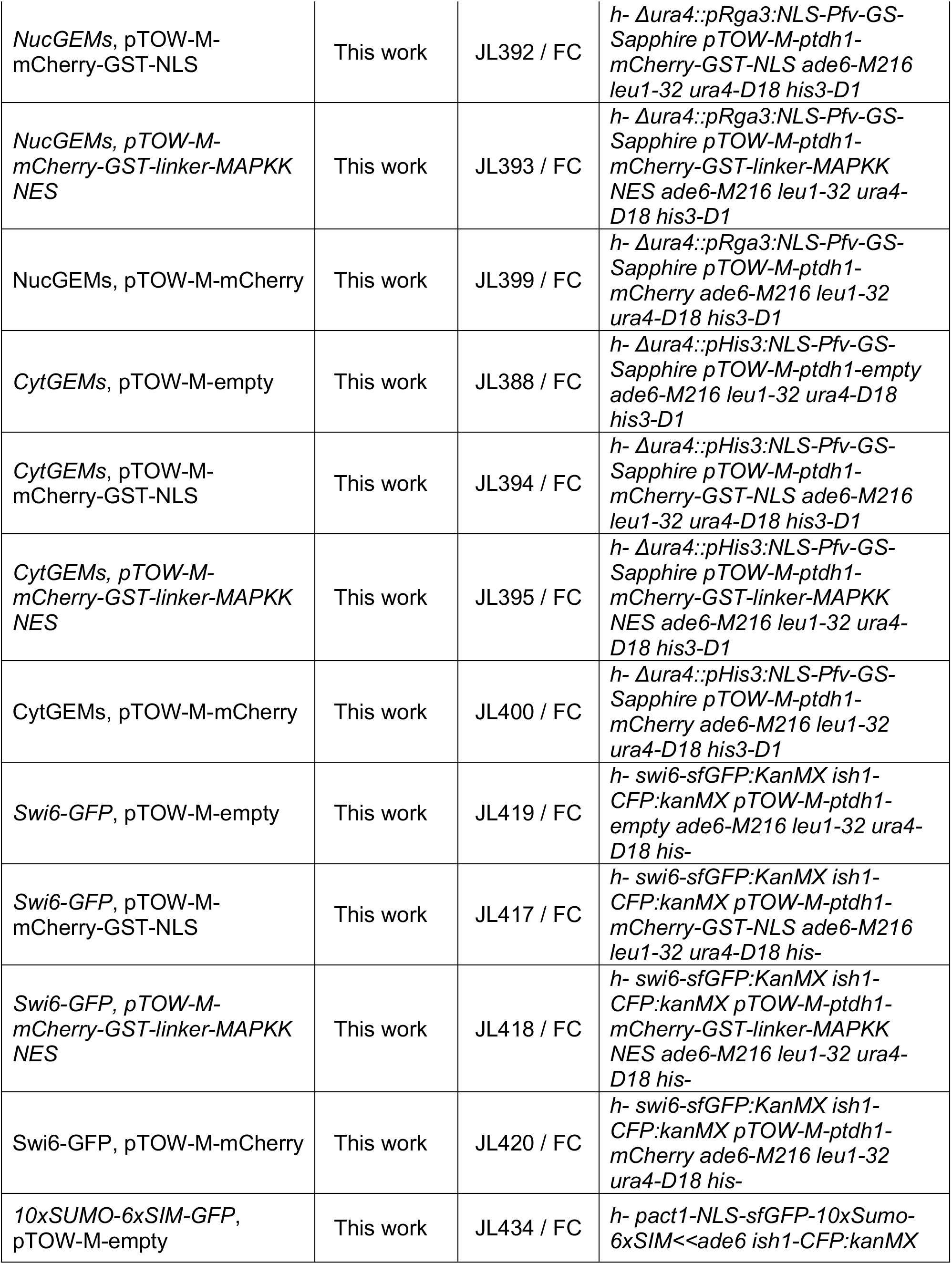
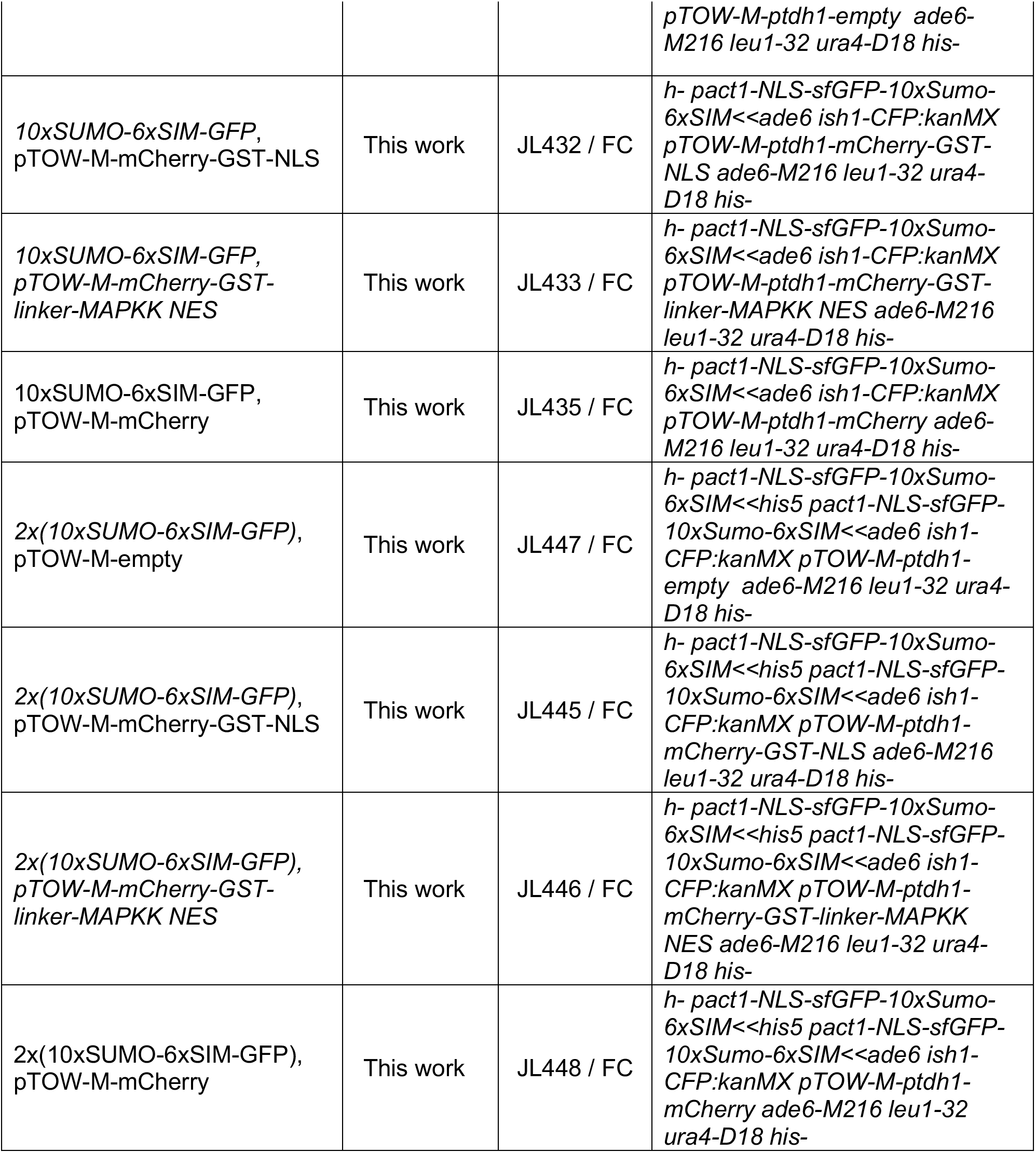

## Supplemental figures

**Supplementary Fig. 1.**
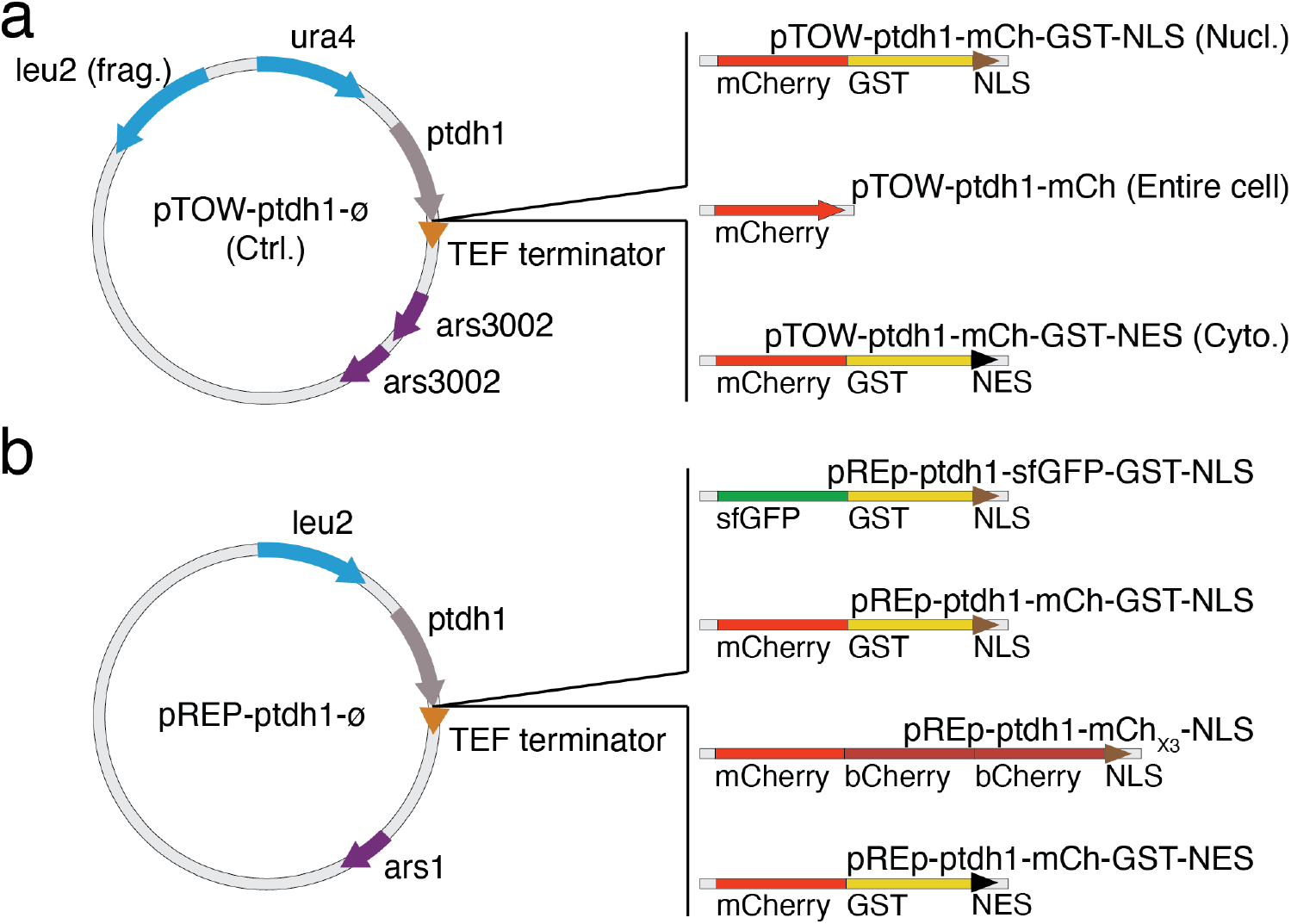
Schematic of plasmid constructs. **a**, Schematic of the multi-copy plasmid construction based on the pTOW-ptdh1 backbone expressing an mCherry-tagged protein. **b**, Examples of different plasmids leading to various protein combination constructs where the promoter, fluorophore and GST were swapped compared to the pTOW backbone in **a**.

**Supplementary Fig. 2.**
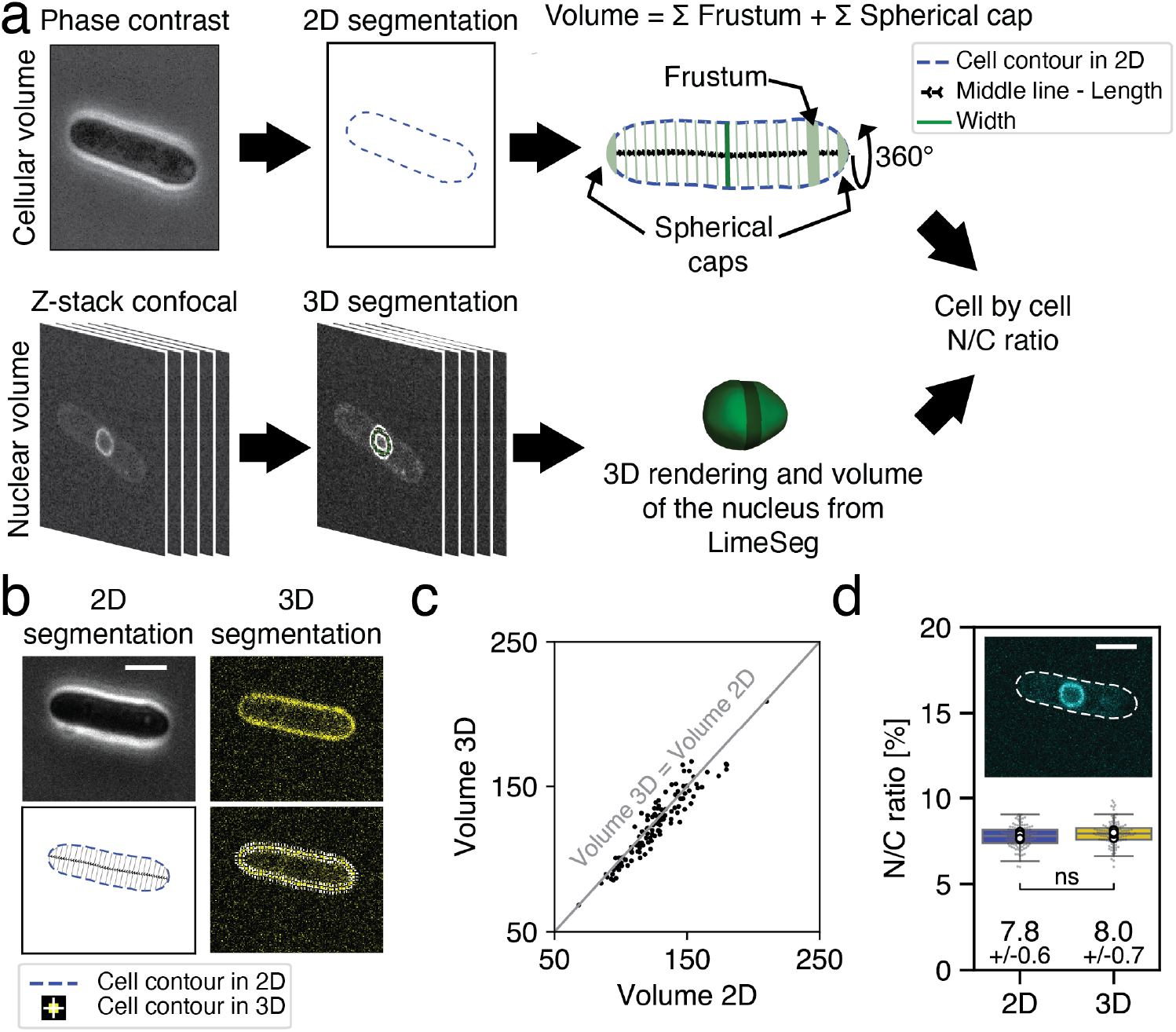
Cellular and nuclear volume measurement pipeline. **a,** Schematic illustrating the methods for measurement of the N/C ratio for individual cells. A phase-contrast image of the mid-plane is used to segment the cell in 2D. Principal component analysis of the cell contour defines the major axis and midline, along which a stack of frustums matching the cell outline is constructed. Skeletonization is used to segment the cell into frustums of equal width, with spherical caps at each cell tip. Assuming rotational symmetry around the midline, the total cell volume is calculated (top). Bottom, nuclear volume determination from a 3D confocal z-stack. The nuclear envelope is segmented using the LimeSeg plugin, yielding a 3D rendering and nuclear volume. **b,** Representative mid-plane phase-contrast and confocal images of the same cell expressing a plasma membrane marker (yellow), used to compare cell volume measurements obtained from 2D segmentation with those obtained from full 3D segmentation using LimeSeg. Scale bar, 5 µm. **c,** Comparison of cellular volumes calculated using the 2D segmentation pipeline and 3D LimeSeg-based segmentation for a population of fission yeast cells (N = 118), showing good agreement between methods. **d,** Comparison of N/C ratio measurements for the same cell population as in (c) obtained using combined 2D cell volume and 3D nuclear volume measurements versus fully 3D segmentation for both cell and nucleus, demonstrating consistent results across methods. Replicate-level paired permutation test comparing N/C ratios between 2D and 3D conditions. Each replicate’s mean was used to account for replicate-to-replicate variability, indicating no significant difference. Mean N/C ratio values and standard deviation.

**Supplementary Fig. 3.**
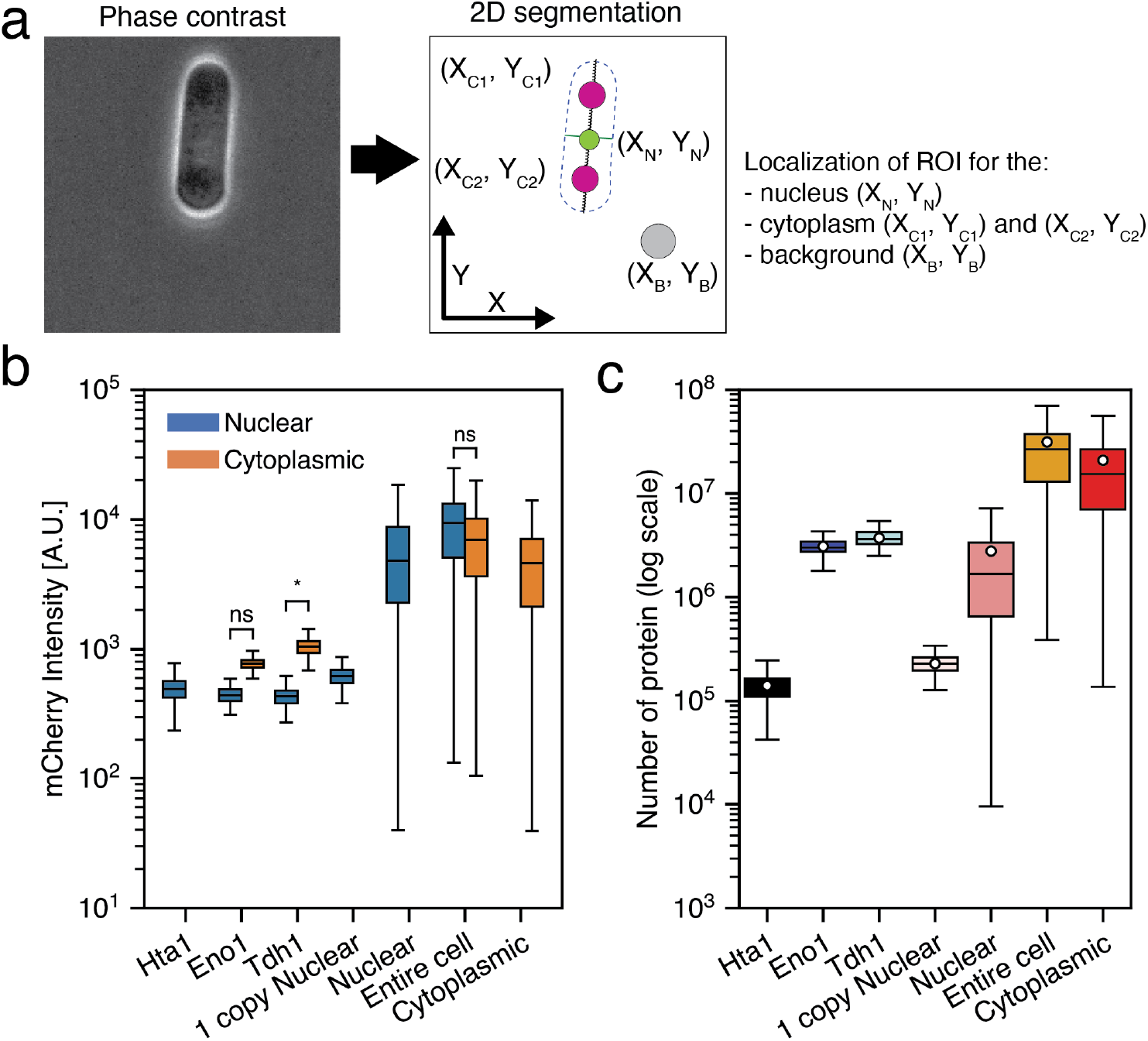
Proteins intensity measurements and quantification analysis. **a,** Representative phase-contrast image of fission yeast cell used for 2D-sgmentation (left), showing the region of interests (ROIs) automatically selected after segmentation to measure fluorescence intensities in the nucleus, cytoplasm and the background (right). Two cytoplasmic ROIs were defined to account for potential asymmetric protein distribution between the tips. **b,** Quantification of mCherry fluorescence intensity for each condition (**see** Fig. 1a), showing the distribution of the signal between the nucleus (blue) and the cytoplasm (gold). When indicated, nuclear and cytoplasmic intensities from the same cells were compared using a two-sided Wilcoxon signed-rank test. Note that the Entire Cell strain shows no significant difference between the nuclear and cytoplasmic mCherry fluorescence intensities (N≥82 cells per conditions) **c,** Quantification of total mCherry protein levels per cells for each condition.

**Supplementary Fig. 4.**
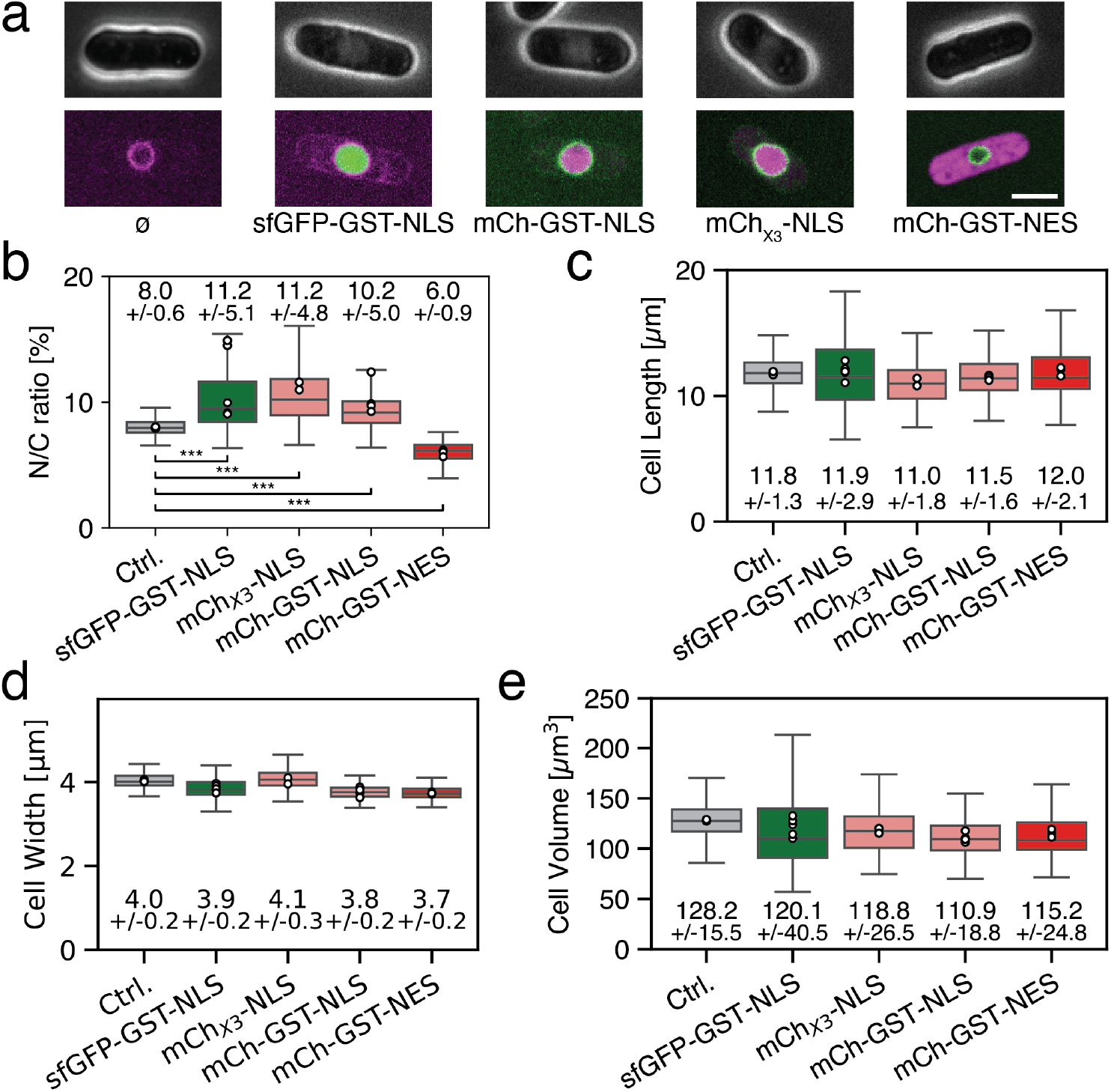
Expression of other proteins besides mCherry-GST also alter the N/C ratio. **a,** Phase contrast images (top row) and mid-plane confocal images (bottom row) of representative cells for each plasmid construct shown in **Supplementary Fig. 1,b**, with the nuclear membrane and the exogenous protein tagged. **b,** Quantification of N/C ratios for populations of cells corresponding to the constructs in **a**, showing that neither mCherry nor GST alone drives changes in nuclear volume. Boxplots show single-cell N/C ratios, with each point representing the mean of an individual biological replicate. Statistical significance relative to the control was assessed using a linear mixed-effects model with strain as a fixed effect and replicate as a random intercept. Significance is indicated below each comparison (N≥162 cells per conditions). **c-e)** Quantification of the cell’s geometrical parameters: cell length **e**, width **f**, and volume **g**. Mean values and standard deviation shown. Scale bar, 5 µm

**Supplementary Fig. 5.**
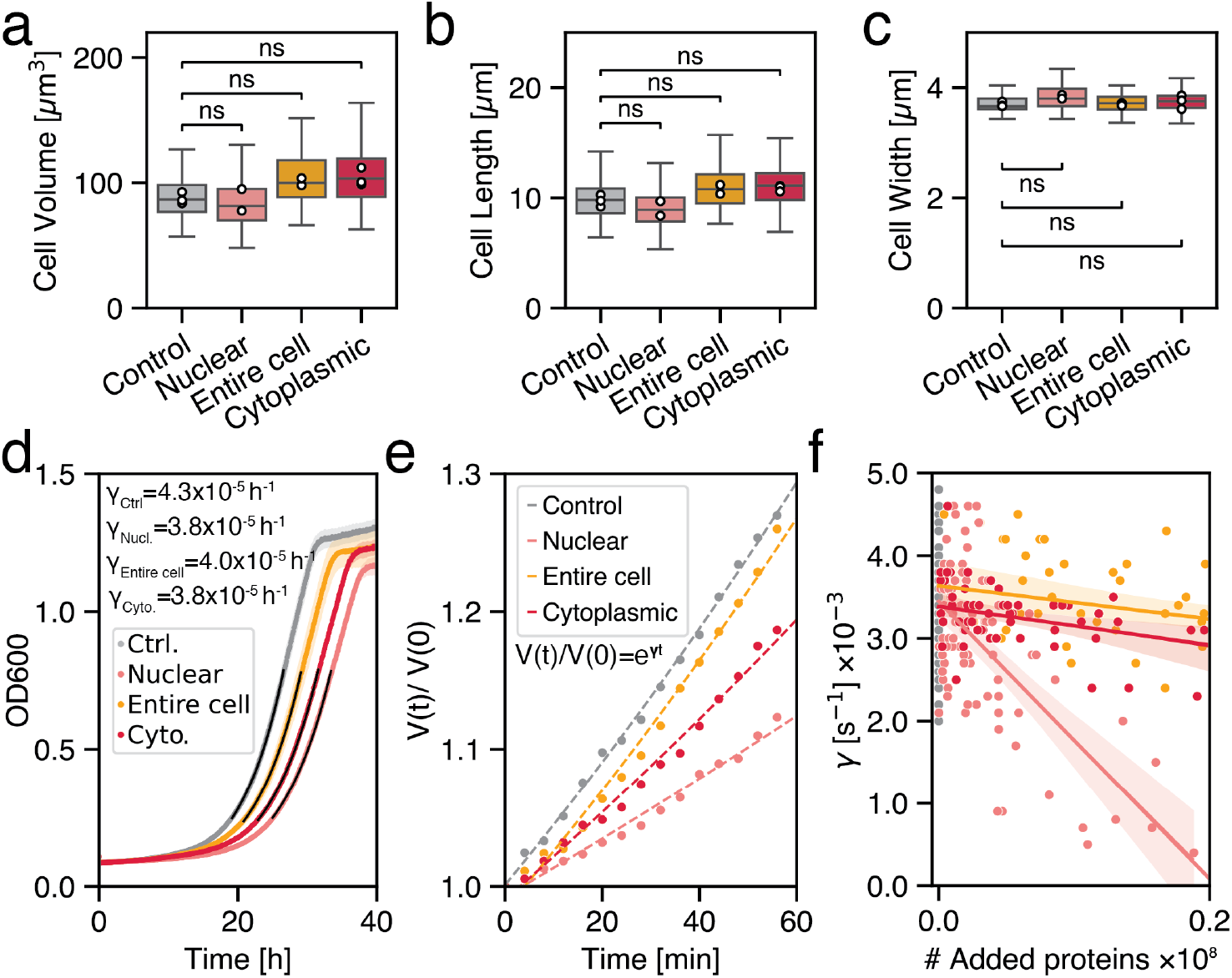
Effects of expression of mCherry-based proteins on cell size and growth rate depend on their localization. **a-c,** Cells width, length and volumes measured using our pipeline for each condition described in Fig.1a. Statistical analysis was performed on replicate means using a one-way, followed by a Welch’s t-test comparing each strain to the control strain (Control) with Bonferroni correction. **d,** Mean growth curves of cells in liquid culture for each condition (N=7 curves per conditions). Optical density at 600 nm (OD600) was measured over time. Population growth rates were estimated by fitting the exponential phase of the curves (black lines). **E,** Representative single-cell normalized volume trajectories over time for each strain. Each single-cell growth curve was fitted with a simple exponential function to obtain a single-cell growth rate (γ). **f,** Single-cell growth rates plotted as a function of the amount of extra protein per cell, for all condition (N>106 cells per condition).

**Supplementary Fig. 6.**
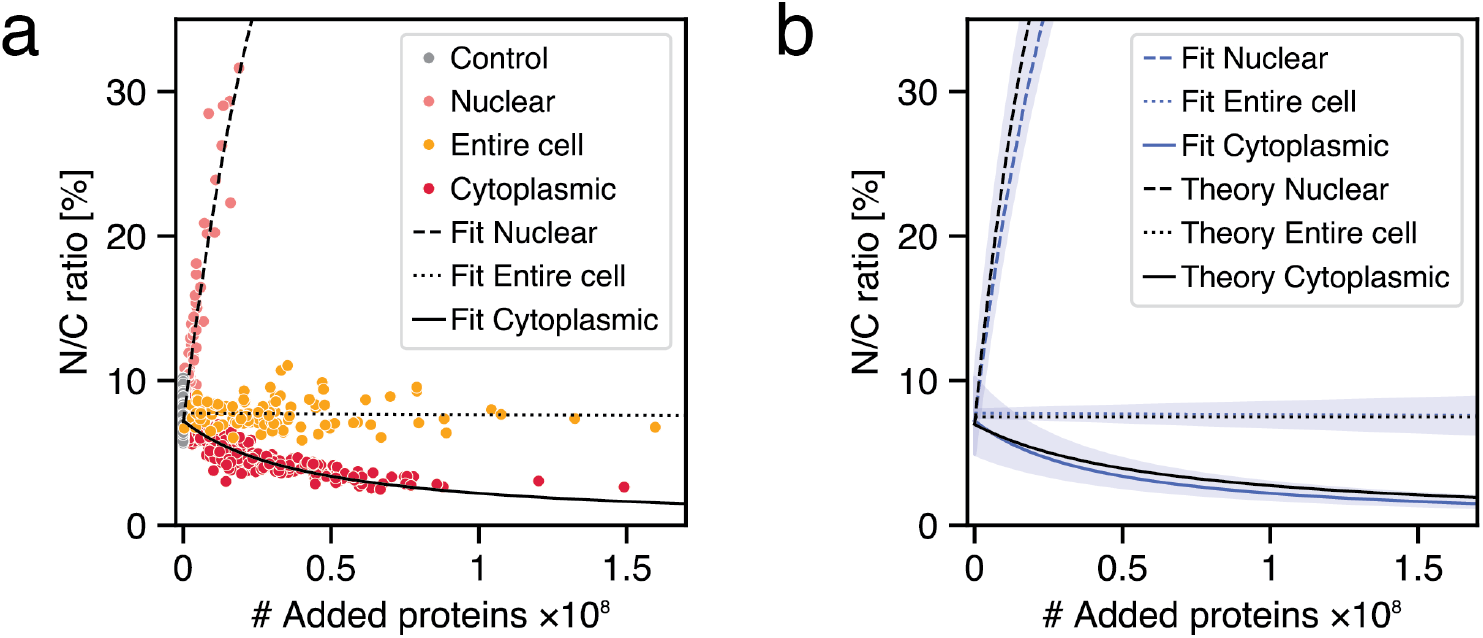
Quantitative agreement between experimental data and theory strongly supports an osmotic model. **a,** Experimental data from Fig. 1d showing the N/C ratio as a function of the amount and subcellular localization of added exogenous protein. Fits to equations (7) and (8) provide estimates for the total protein quantity in the nucleus and the cell. **b,** Comparison between fit curves from **a** and theoretical curves, for each condition. Shaded regions in blue indicate the 1σ uncertainty of the fit curves in **a**. Theoretical curves (from Fig. 1d) are based upon a total protein number from proteomics data, which predict the expected behavior for the N/C ratio in each condition.

**Supplementary Fig. 7.**
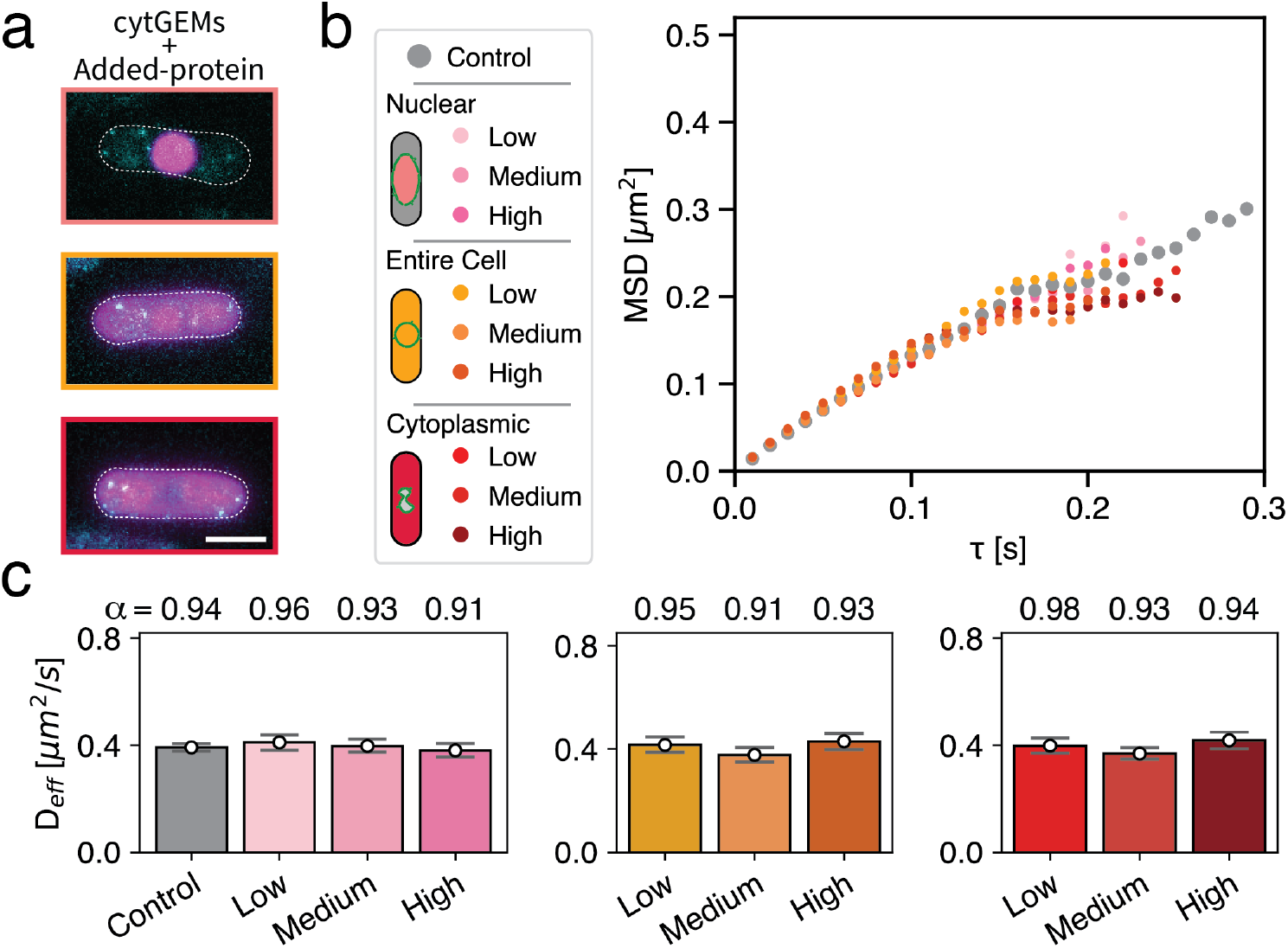
Expression of mCherry-based proteins does not affect cytoplasmic mesoscale diffusivity. **a,** Representative fluorescence images of fission yeast expressing cytoplasmic GEMs (cyan) and the added protein tagged with mCherry (magenta). Scale bar 5 µm. **b,** Mean squared displacement (MSD) of cytoplasmic GEMs for each condition. Except for the control, the population was split into low, medium, and high levels of mCherry expression. **c,** Effective diffusion coefficient extracted from the GEM movements in each condition, plotted as a function of added protein expression level. Anomalous exponent (α) are indicated above each bar.

**Supplementary Fig. 8.**
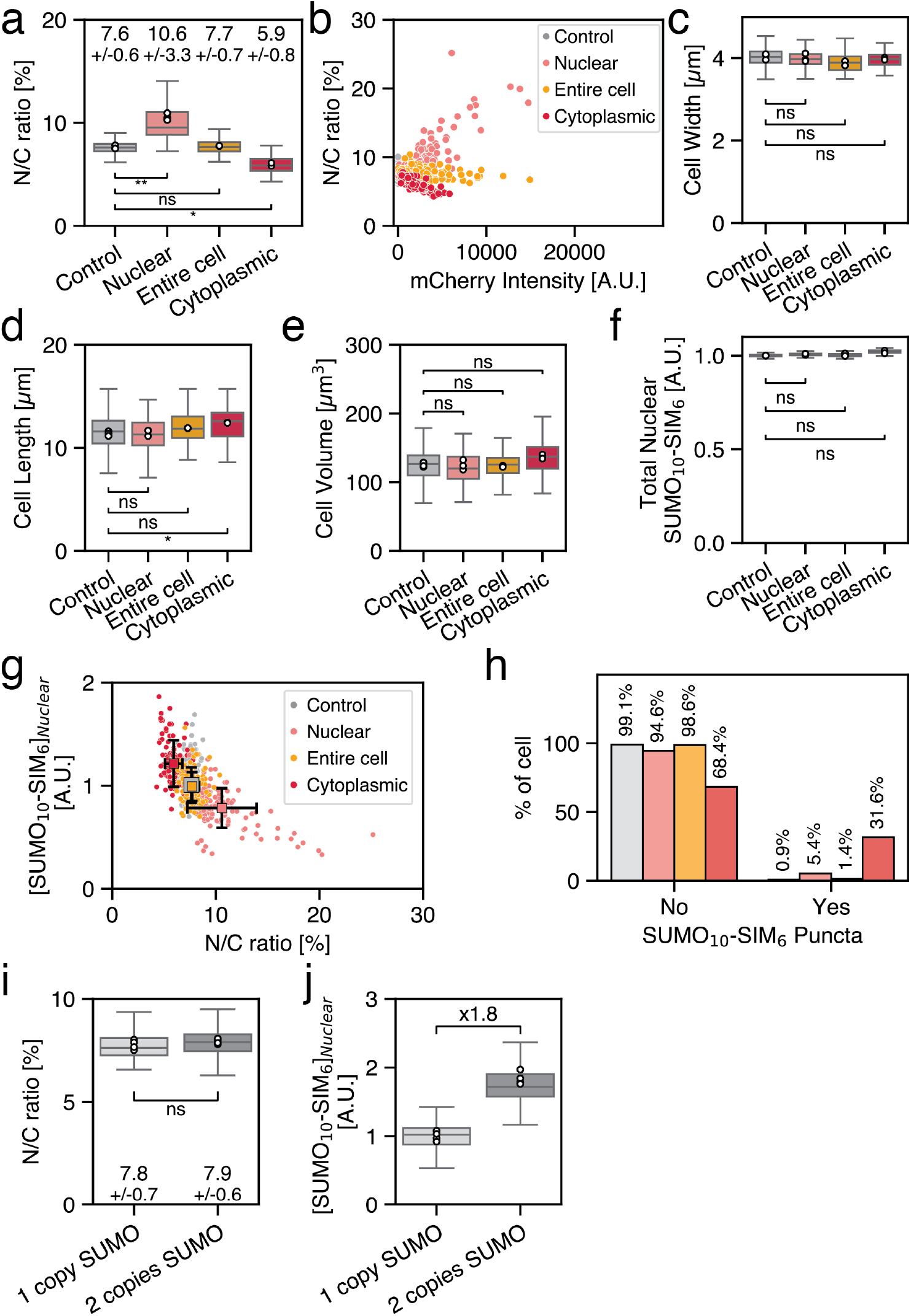
Characterization of cells carrying one copy of the synthetic condensate gene. **a,** N/C ratio measured in populations of cells carrying one copy of the SUMO_10_-SIM_6_ synthetic condensate gene, expressing additional exogenous proteins (N≥98 cells per condition). Mean values and standard deviations are shown. **b,** N/C ratio plotted as a function of mCherry intensity for each condition in the one copy SUMO_10_-SIM_6_ background. **c-e,** Cells width, length and volume for cells carrying one copy of the SUMO_10_-SIM_6_, measured for each condition using an automated analysis pipeline. **f,** Total nuclear SUMO_10_-SIM_6_ concentration measured for each condition. **g,** Same as **f**, plotted for individual cells and mean values overlaid. **h,** Percentage of cells for which a nuclear SUMO_10_-SIM_6_ punctum was detected in each condition (N≥129 cells per condition). **i,** N/C ratio for populations of cells carrying one or two copies of the SUMO_10_-SIM_6_ construct, showing no difference in the N/C ratio between background. **j,** Nuclear concentration of SUMO_10_-SIM_6_ for the one- and two-copy strain backgrounds. Statistical analyses were performed on replicate means (white dots) using a one-way ANOVA, followed by pairwise Welch’s t-test comparing each strain to the control.

**Supplementary Fig. 9.**
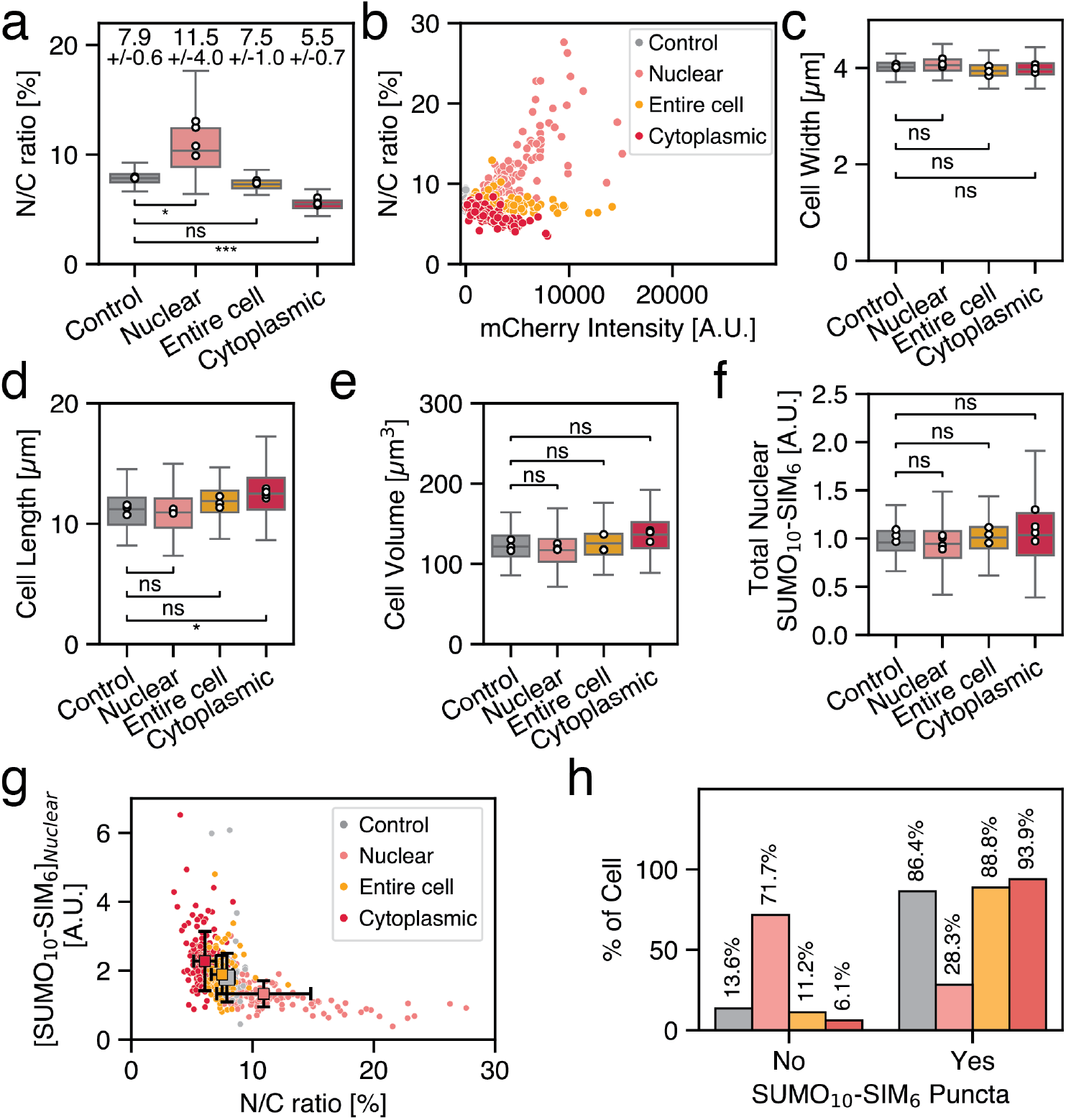
Characterization of cells carrying two copies of the synthetic condensate gene. **a,** N/C ratio measured in populations of cells carrying two copies of the SUMO_10_-SIM_6_ synthetic condensate gene, expressing additional exogenous proteins (N≥107 cells per condition). Mean values and standard deviations are shown. **b,** N/C ratio plotted as a function of mCherry intensity for each condition in the two-copy SUMO_10_-SIM_6_ background. **c-e,** Cells width, length and volume for cells carrying two copies of the SUMO_10_-SIM_6_, measured for each condition using an automated analysis pipeline. **f,** Total nuclear SUMO_10_-SIM_6_ concentration measured for each condition. **g,** Same as **f**, normalized by the intensity of the 1 copy SUMO_10_-SIM_6_ background and plotted for individual cells and mean values overlaid. **h,** Percentage of cells for which a nuclear SUMO_10_-SIM_6_ punctum was detected in each condition (N≥103 cells per condition). Statistical analyses were performed on replicate means (white dots) using a one-way ANOVA, followed by pairwise Welch’s t-test comparing each strain to the control.

**Supplementary Fig. 10.**
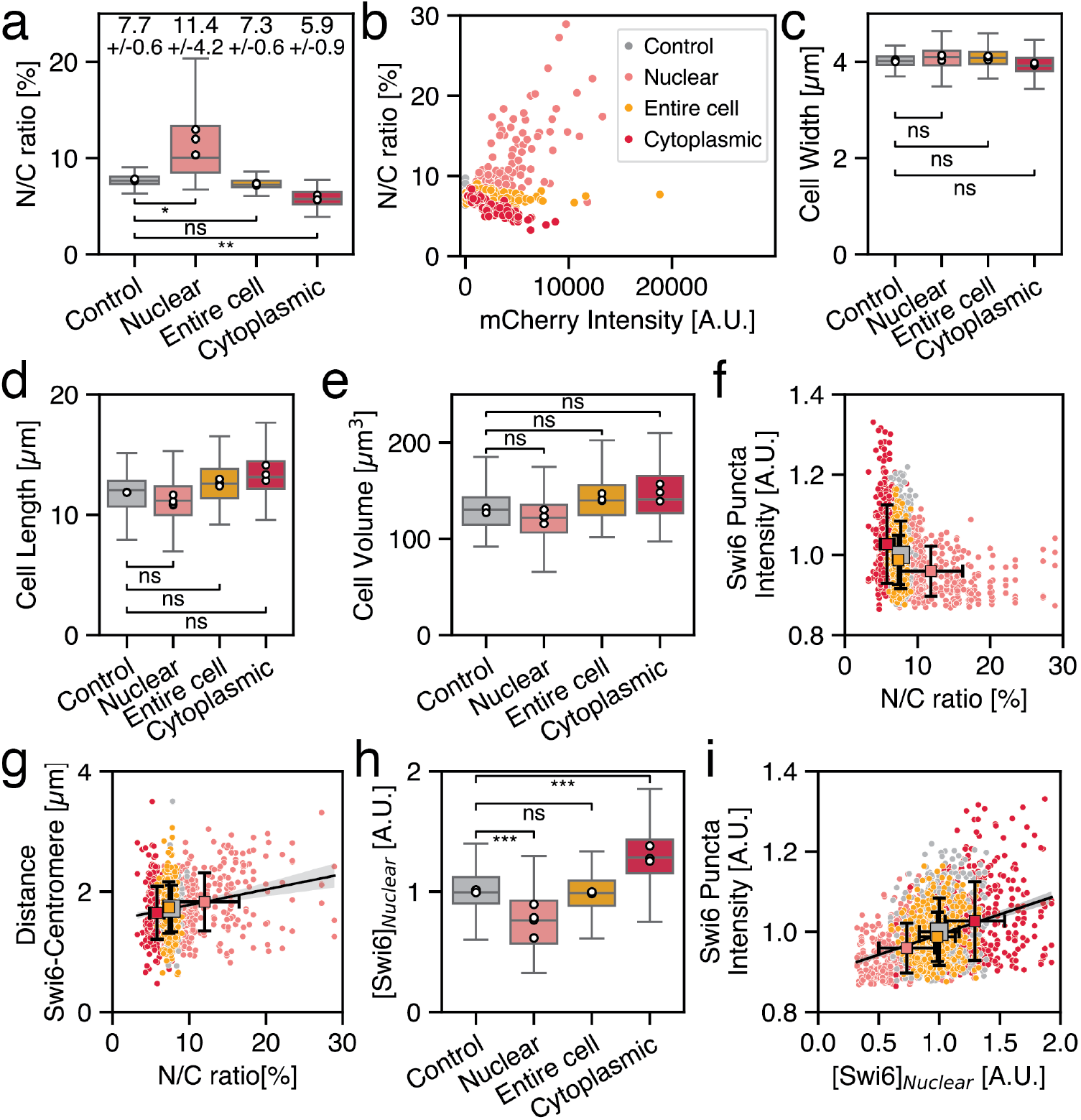
Effect of nuclear size on Swi6-GFP puncta intensity and nuclear concentration. **a,** N/C ratio measured in populations of cells expressing additional exogenous proteins and carrying Swi6-tagged in their background (N≥188 cells per condition). Mean values and standard deviations are shown. **b,** N/C ratio plotted as a function of mCherry intensity for each condition. **c-e)** Cells width, length and volume measured for each condition. **f,** Swi6-GFP puncta intensity as a function of the N/C ratio, with means and standard deviation overlaid (N=1741 puncta). **g,** Distance between Swi6-GFP non-centromere puncta and the centromere as a function of the N/C ratio (black line indicates a linear fit with a positive slope; N=1741 puncta). **h,** Nuclear concentration of Swi6-GFP for each condition. **i,** Swi6-GFP puncta intensity plotted as a function of the nuclear concentration of Swi6-GFP. Means and standard deviation are overlaid. Statistical analyses were performed on replicate means (white dots) using one-way ANOVA tests (a,c-e,h).

**Supplementary Fig. 11.**
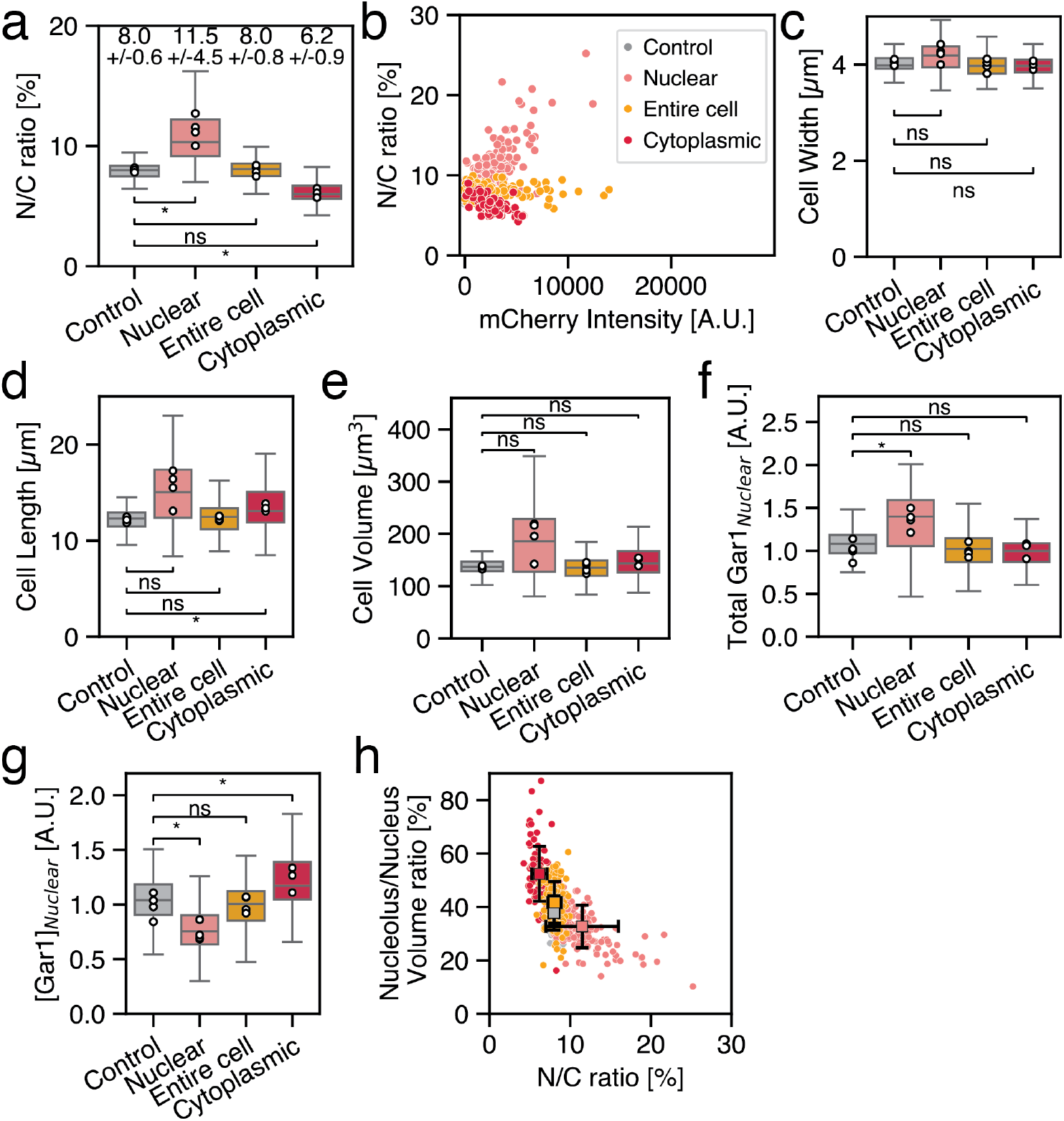
Effects of nuclear size on the nucleolar marker Gar1-GFP. **a,** N/C ratio measured in populations of cells expressing additional exogenous proteins and carrying endogenously tagged Gar1 (N≥97 cells per condition). Mean values and standard deviations are shown. **b,** N/C ratio as a function of mCherry fluorescence intensity for each condition. **c-e)** Cells width, length and volume measured for each condition. **f,** Total nuclear Gar1-GFP signal for each condition **g,** Nuclear concentration of Gar1-GFP for each condition. **h,** Nucleolus to nucleus volume ratio as a function of the N/C ratio plotted per cell, with means and standard deviation overlaid for each condition. Statistical analyses were performed on replicate means (white dots) using one-way ANOVA (a,c-g).

